# LFQ Benchmark Dataset - Generation Beta: Assessing Modern Proteomics Instruments and Acquisition Workflows with High-Throughput LC Gradients

**DOI:** 10.64898/2026.01.29.702266

**Authors:** Bart Van Puyvelde, Robbe Devreese, Cristina Chiva, Eduard Sabidó, Sibylle Pfammatter, Christian Panse, Jeewan Babu Rijal, Charline Keller, Ihor Batruch, Patrick Pribil, Jean-Baptiste Vincendet, Frédéric Fontaine, Lars Lefever, Pedro Magalhães, Dieter Deforce, Paolo Nanni, Bart Ghesquière, Yasset Perez-Riverol, Lennart Martens, Christine Carapito, Robbin Bouwmeester, Maarten Dhaenens

**Affiliations:** ProGenTomics, Laboratory of Pharmaceutical Biotechnology, Ghent University, 9000 Ghent, Belgium; VIB-UGent Center for Medical Biotechnology, VIB, 9052 Ghent, Belgium; Department of Biomolecular Medicine, Ghent University, 9052 Ghent, Belgium; Centre for Genomic Regulation, The Barcelona Institute of Science and Technology (BIST), Dr Aiguader 88, 08003 Barcelona, Spain; Univeristat Pompeu Fabra, Dr. Aiguader 88, 08003 Barcelona, Spain; Functional Genomics Center Zurich (FGCZ) - University of Zurich/ETH Zurich, Winterthurerstrasse 190, CH-8057 Zurich, Switzerland; Swiss Institute of Bioinformatics (SIB), Quartier Sorge - Batiment Amphipole, CH-1015 Lausanne, Switzerland; BioOrganic Mass Spectrometry Laboratory (LSMBO), IPHC UMR 7178, University of Strasbourg, CNRS, 67087 Strasbourg, France; Infrastructure Nationale de Protéomique ProFI – UAR 2048, Strasbourg, 67087, France; Sciex, Concord, Ontario, Canada; Thermo Fisher Scientific, Paris, France; Laboratory of Applied Mass Spectrometry, Department of Cellular and Molecular Medicine, KU Leuven, Leuven, 3000, Belgium; Metabolomics Expertise Center, Center for Cancer Biology VIB, Leuven, 3000, Belgium; VIB-KU Leuven Center for Brain & Disease Research Technologies, Proteomics Unit, Leuven, Belgium; European Molecular Biology Laboratory, European Bioinformatics Institute, Wellcome Genome Campus, Hinxton, UK

## Abstract

Recent advances in liquid chromatography–mass spectrometry (LC-MS) have accelerated the adoption of high-throughput workflows that deliver deep proteome coverage using minimal sample amounts. This trend is largely driven by clinical and single-cell proteomics, where sensitivity and reproducibility are essential. Here, we extend our previous benchmark dataset (PXD028735) using next-generation LC-MS platforms optimized for rapid proteome analysis. We generated an extensive DDA/DIA dataset using a human-yeast-*E. coli* hybrid proteome. The proteome sample was distributed across multiple laboratories together with standardized analytical protocols specifying two short LC gradients (5 and 15 min) and low sample input amounts. This dataset includes data acquired on four different platforms, and features new scanning quadrupole-based implementations, extending coverage across different instruments and acquisition strategies. Our comprehensive evaluation highlights how technological advances and reduced LC gradients may affect proteome depth, quantitative precision, and cross-instrument consistency. The release of this benchmark dataset via ProteomeXchange (PXD070049 and PXD071205), allows for the acceleration of cross-platform algorithm development, enhance data mining strategies, and supports standardization of short-gradient, high-throughput LC-MS-based proteomics.

## Background & Summary

Proteomics has rapidly evolved into a cornerstone of molecular biology, providing unparalleled insight into protein function, interactions, and regulation. While DNA and RNA measurements reveal what a cell is capable of, proteomics captures what it is actually doing—reflecting the dynamic protein complement and post-translational modifications (PTMs) that define cellular function and phenotype.

The complexity of the proteome, shaped by alternative splicing, variable translation rates, and diverse PTMs, necessitates analytical approaches capable of resolving this intricate molecular landscape with both depth and precision. Liquid Chromatography coupled to Mass spectrometry (LC-MS)-based proteomics has emerged as the leading technology for large-scale protein analysis, offering an unbiased approach to protein identification and quantification ^1^. Advancements in mass spectrometry instrumentation, including Zeno trapping, next-generation time-of-flight (TOF) analyzers such as the multi-reflectron TOF, and emerging hybrid platforms like the Orbitrap Astral, have significantly enhanced sensitivity, acquisition speed, and dynamic range ^2–6^. These advancements have propelled proteomics from bulk tissue analysis toward increasingly challenging applications, such as clinical proteomics, single-cell proteomics and spatially resolved protein mapping ^7^.

While data-dependent acquisition (DDA) has been the cornerstone for many years, it has now been taken over by data-independent acquisition (DIA) due to its increased reproducibility and sensitivity ^8,9^. DDA relies on sequential selection of precursors for fragmentation and identification, but this approach is inherently limited by sampling bias. In contrast, DIA captures all fragment ions across a defined mass range, overcoming selection bias and enabling more comprehensive proteome coverage. These gains come with a trade-off, as DIA generates complex data that is difficult to interpret. In response, the field has rapidly developed advanced computational workflows capable of handling these complexities ^10^. Machine learning has rapidly emerged as a powerful method to predict the behavior of peptides, including their fragmentation spectra, retention times and collisional cross-sections ^11–18^. These predicted properties can then be incorporated into spectral libraries to expand coverage and improve the accuracy of peptide identification and quantification ^19,20^.

Additional separation dimensions, such as ion mobility, like TIMS and FAIMS, have been introduced to enhance resolution and reduce spectral overlap ^21–24^. Alternatively, some have implemented scanning quadrupole based acquisition, i.e. SONAR ^25^, Scanning SWATH ^26^ or ZT Scan DIA ^27^ and diagonalPASEF ^28^ (midia-PASEF ^29^ and SynchroPASEF ^30^), providing an extra dimension to alleviate the spectral complexity by sampling a sliding quadrupole selection window with a very fast TOF detector. Others have implemented the usage of narrow-window DIA (nDIA) to increase the specificity ^6^.

A key driver behind these innovations is the growing field of single-cell proteomics, wherein the objective is to profile the proteome at the resolution of a single cell. This field specifically must address the technical challenges associated with analyzing ultra-low amounts of material ^31^. The demand for measuring thousands of proteins from a single cell has pushed instruments toward higher speed and sensitivity ^32^. To meet the high-throughput requirements of single-cell studies, workflows increasingly use extremely short LC gradients. By minimizing sample carryover and leveraging low flow rates, these ultra-fast chromatographic workflows allow simultaneous detection of a large number of proteins from sub-nanogram sample inputs. During method development, many groups work with “single-cell equivalent” inputs, commonly ∼250 pg, to emulate the average protein content of a single cell ^33^. These developments made to meet the demands of single-cell proteomics underscore the importance of combining accelerated LC separations, minimal sample requirements, and advanced DIA ^34,35^.

The shift from DDA to DIA, together with increasingly complex acquisition strategies, has expanded the analytical possibilities of proteomics, yet equally magnifies the need for standardized benchmarks. In particular, the rise of single-cell proteomics has reinforced the importance of shared datasets that enable consistent evaluation of workflows across diverse experimental contexts. Our previous LFQ benchmark study (PXD028735) has become a widely used community resource, with 145,147 PRIDE downloads since 2022, placing it within the top 1% of PRIDE datasets by downloads. Therefore, we present a second-generation LFQ benchmark study that builds upon our previous dataset by evaluating the performance of the latest generation MS instruments, i.e. SCIEX ZenoTOF 7600+ system & ZenoTOF 8600 system, Bruker Ultra 2, and Thermo Orbitrap Astral, on recent DIA, narrow-window DIA, and scanning quadrupole DIA methodologies (diagonal-PASEF, ZT Scan DIA) under conditions relevant to bulk as well as single-cell proteomics ^36^. We adapt the widely used benchmark experimental design employing a triple proteome-mixture ^37^, and include adequate technical replicates in both nano and capillary flow LC setups. In line with our previous benchmark, our aim is not to maximize peptide and protein identifications but to create a comprehensive, community-driven resource for the proteomics field. This dataset is intended to facilitate transparent comparisons of algorithms and workflows and already contributes to the ongoing development of community initiatives such as ProteoBench ^38^, helping to shift the focus beyond simple metrics of identification counts toward more rigorous assessment of quantitative accuracy and precision.

## Methods

### Sample preparation

Mass spectrometry-compatible protein digest extracts from Human K562 (P/N: V6951, Lot: 0000608712) and Yeast (P/N: V7461, Lot: 0000601330) were obtained from Promega (Madison, Wisconsin, USA), while the lyophilized MassPrep *Escherichia coli* protein digest extract (P/N: 186003196, Lot: W01111913) was sourced from Waters Corporation (Milford, Massachusetts, USA). Each extract underwent reduction with dithiothreitol (DTT), alkylation with iodoacetamide (IAA), and digestion with sequencing grade Trypsin (-Lys C) by the respective manufacturers. The digested protein extracts were reconstituted in a solution of 0.1% formic acid (FA) in 5% acetonitrile (Biosolve B.V, Valkenswaard, The Netherlands). Following the approach of Navarro et al.^37^, two master samples, A and B, were prepared as illustrated in Figure 1. Sample A consisted of Human, Yeast, and *E. coli* digests mixed at 65%, 30%, and 5% weight/weight (w/w), respectively. Sample B was formulated by combining Human, Yeast, and *E. coli* protein digests at 65%, 15%, and 20% w/w, respectively. Additionally, a third mixture, Sample C, was created with Human, Yeast, and *E. coli* protein digests at 65%, 3%, and 32% w/w, respectively, to assess the impact of larger ratio differences on analytical performance. Sample injection order was performed using a standardized scheme (see Supplementary Table 1), in which replicates of the same sample were intentionally interspersed rather than measured consecutively to minimize batch effects.

**Figure 1.**
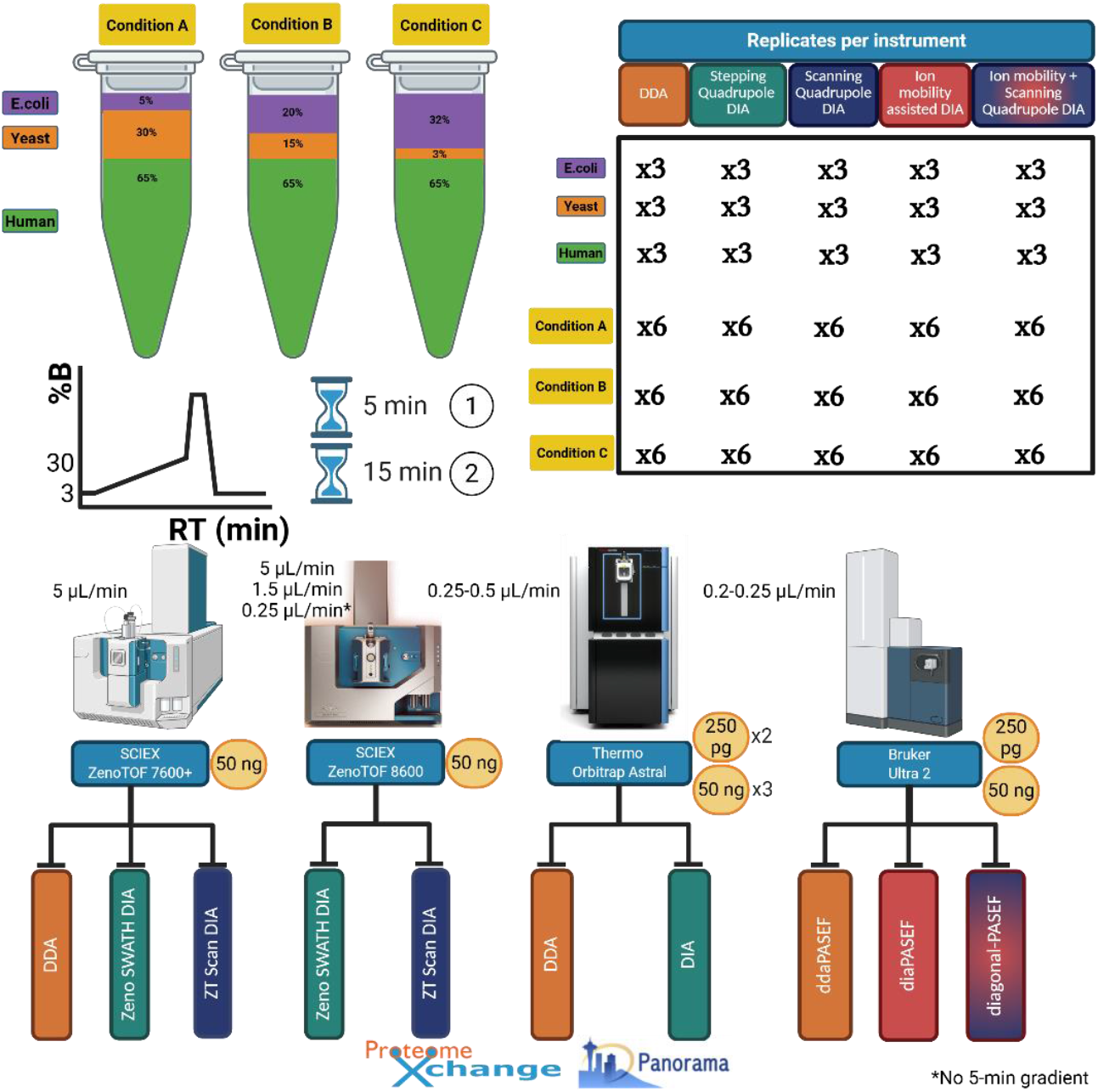
Schematic overview of Acquisition Strategies and Instrumentation. A comprehensive LC-MS dataset was generated using commercial Human K562, Yeast, and *Escherichia coli* (*E. coli*) proteome digests. To simulate experimental ratio differences, three hybrid proteome samples (A, B, and C) containing defined proportions of tryptic peptides from Human, Yeast, and *E. coli* were prepared. Both the individual proteome digests and the hybrid mixtures were analyzed using two short LC gradient lengths (5- and 15-minutes active separation) to assess the impact of chromatographic speed on data quality and proteome coverage. Data acquisition was performed in both Data-Dependent Acquisition (DDA) and Data-Independent Acquisition (DIA) modes across four advanced LC-MS platforms: SCIEX ZenoTOF 7600+ and 8600, Thermo Scientific Orbitrap Astral (2 sites), and Bruker timsTOF Ultra 2. Each commercial lysate was measured individually in triplicate, while the hybrid mixtures were analyzed in six replicates per gradient and acquisition method, ensuring robust statistical comparisons. The complete dataset is publicly available via ProteomeXchange under the identifiers PXD070049 and PXD071205, providing a valuable resource for the proteomics community. Additionally, system suitability monitoring was integrated into the workflow using commercial *E. coli* lysate digest. These QC samples were acquired in DDA at multiple time points throughout the batch on each instrument. The AutoQC workflow automatically imported QC data into Skyline and uploaded it to the Panorama AutoQC server via AutoQC Loader, enabling real-time system suitability assessment across all LC-MS platforms used in the study. Figure created in BioRender.

### LC-MS

In this section, a detailed description of the different LC-MS/MS parameters is given for each instrumental setup applied to generate this comprehensive dataset. Most of the instruments were operated according to the lab’s best practice, i.e. not necessarily the best attainable, but rather most realistic data quality. A second Orbitrap Astral dataset was generated under more refined conditions. The 1.5 µL/min and nano-flow dataset on the ZenoTOF 8600 was generated by the SCIEX demo lab. The sample load was kept consistent across each lab at 50 ng per injection, which is 100-fold lower compared to our previous LFQ benchmark dataset ^36^. Additionally, some laboratories measured 250 pg injections to mimic single-cell conditions. Samples were analyzed using two different standard gradients which included a 5-min and a 15-min active separation ramp of 3-30% acetonitrile content as a set criteria followed to the best efforts, with the rest of the method adjusted by the operators to accommodate LCs and workflows variety. Also, acquisitions and results were not meant to either reflect or conflict with the literature. Differences in the absolute number of identified peptides and proteins can be attributed to factors such as LC conditions (flow rate, column), MS instrumentation, operator choices (instrument settings e.g. window size), and search algorithm compatibility; therefore, direct conclusions on MS instrument performance cannot be drawn from this dataset.

### SCIEX ZenoTOF 7600+ system (Capillary flow LC – 5 µL/min) – 50 ng

An ACQUITY™ M-Class™ UPLC System (Waters Corporation, Milford, MA) was used and operated in direct injection mode with the SCIEX ZenoTOF 7600+ system (SCIEX, Concord, Ontario, Canada). Mobile phases consisted of 0.1% FA in water (A) and 0.1% FA in acetonitrile (B). Microflow separations were done using a Kinetex XB-C18 (15 cm x 300 µm, 2.6 µm particle size)(Phenomenex, California, USA) column at a flow rate of 5 µL/min and column temperature of 30 °C using the following gradient: i) after an initial hold at 3% B for 1 min, a gradient from 3% to 30% mobile phase B in 5 or 15 min, respectively, was applied ii) ramp to 80% mobile phase B in 1 min. The washing step at 80% mobile phase B lasted 2 min and was followed by an equilibration step at 3% mobile phase B (starting conditions) for 7 min. An OptiFlow Turbo V ion source was used with a microflow probe (1-10 µL/min electrode) for the microflow experiments, with source parameters as following: GS1: 20 psi, GS2: 60 psi, Curtain Gas: 35 psi, CAD gas: 7 psi, source temperature: 200 °C and Ionspray voltage: 5000V.

#### Data-Dependent Acquisition

For the 5-min gradient (a cycle time of 0.573 s), MS1 spectra were collected between 400-1200 m/z for 100 ms. The 30 most intense precursors ions with charge states 2-4 that exceeded 300 counts per second were selected for fragmentation, and the corresponding fragmentation MS2 spectra were collected between 140-1750 m/z for 10 ms. After the fragmentation event, the precursor ions were dynamically excluded from reselection for 3 s.

For the 15-min gradient (a cycle time of 0.795 s), MS1 spectra were collected between 400-900 m/z for 100 ms. The 45 most intense precursors ions with charge states 2-4 that exceeded 300 counts per second were selected for fragmentation, and the corresponding fragmentation MS2 spectra were collected between 140-1800 m/z for 10 ms. After the fragmentation event, the precursor ions were dynamically excluded from reselection for 4 s.

#### Zeno SWATH DIA

The 5 min-Zeno SWATH DIA experiments used 60 variable windows (Supplementary Table 2) spanning the TOF MS mass range 400-900 Da and MS/MS mass range 140-1800 Da, with Zeno trap pulsing turned on, with MS/MS accumulation times of 7 msec. Before each Zeno SWATH DIA MS cycle an additional MS1 survey scan from 400-900 Da was recorded for 50 ms.

The 15min-Zeno SWATH DIA experiments used 65 variable windows (Supplementary Table 3) spanning the TOF MS mass range 400-900 Da and MS/MS mass range 140-1800 Da, with Zeno trap pulsing turned on, with MS/MS accumulation times of 13 msec. Before each Zeno SWATH DIA MS cycle an additional MS1 survey scan from 400-900 Da was recorded for 50 ms.

#### ZT Scan DIA

For the 5-min ZT Scan DIA experiments, 5 Da Q1 isolation windows were scanned at 750 Da/s to accommodate PWHH values ≤1 s, resulting in an MS2 accumulation time of 6.7 ms. For the 15-min ZT Scan DIA experiments, 10 Da Q1 isolation windows were scanned at 750 Da/s to accommodate PWHH values between 1.1 and 4.5 s, resulting in an MS2 accumulation time of 13.3 ms. For both gradients, the TOF MS and MS/MS mass range was set to 400-900 Da and 140-1750 Da, respectively.

In Sciex OS 3.4, the acquired ZT Scan DIA raw data is calibrated using residual precursor signals and converted to SCIEX’s proprietary .wiff and .wiff.scan file formats.

### SCIEX ZenoTOF 8600 system – 50 ng

An ACQUITY™ M-Class™ UPLC System (Waters Corporation, Milford, MA) was used and operated in direct injection mode with the SCIEX ZenoTOF 8600 system (SCIEX, Concord, Ontario, Canada). Mobile phases consisted of 0.1% FA in water (A) and 0.1% FA in acetonitrile (B).

### 5 µL/min

The 5 µL/min separations were done using a Kinetex XB-C18 (15 cm x 300 µm, 2.6 µm particle size)(Phenomenex, California, USA) at a column temperature of 35 °C using the following gradient: i) after an initial hold at 3% B for 1 min, a gradient from 3% to 30% mobile phase B in 5 or 15 min, respectively, was applied ii) ramp to 80% mobile phase B in 1 min. The washing step at 80% mobile phase B lasted 2 min and was followed by an equilibration step at 3% mobile phase B (starting conditions) for 7 min.

An OptiFlow Pro ion source was used with a microflow probe (1-10 µL/min electrode) for the microflow experiments, with source parameters as following: GS1: 20 psi, GS2: 60 psi, Curtain Gas: 35 psi, CAD gas: 7 psi, source temperature: 200 °C and Ionspray voltage: 5000V.

### 1.5 µL/min

The 1.5 µL/min separations were done using an Evosep EV1109 (8 cm x 150 µm, 1.5 µm particle size)(Evosep, Odense, Denmark) column at a column temperature of 40 °C using the following gradient: i) after an initial hold at 1% B for 4 min, a gradient from 1% to 30% mobile phase B in 11.5 min, respectively, was applied ii) ramp to 80% mobile phase B in 0.5 min. The washing step at 80% mobile phase B lasted 2 min and was followed by an equilibration step at 1% mobile phase B (starting conditions) for 2 min.

An OptiFlow Pro ion source was used with a microflow probe (1-10 µL/min electrode), with source parameters as following: GS1: 10 psi, GS2: 20 psi, Curtain Gas: 35 psi, CAD gas: 7 psi, source temperature: 100 °C and Ionspray voltage: 5000V.

### 0.250 µL/min – Refined conditions

The 0.250 µL/min separations were done using an Aurora® XS Ultimate column (25 cm x 75 µm, 1.7 µm particle size)(IonOpticks, Victoria, Australia) at a column temperature of 50 °C using a 15- and 30-min gradient: i) after an initial hold at 3% B for 18 min, a gradient from 3% to 35% mobile phase B in 15 or 30 min, respectively, was applied ii) ramp to 65% mobile phase B in 2 min, followed by ramping to 80% in 1 min. The washing step at 80% mobile phase B lasted 4 min and was followed by an equilibration step at 1% mobile phase B (starting conditions) for 15 min.

An OptiFlow Pro Nano ion source was used with a NanoCal probe (<1 µL electrode), with source parameters as following: GS1: 10 psi, Nano cel temperature: 250 °C and Nano spray voltage: 2100V.

#### Zeno SWATH DIA

The Zeno SWATH DIA experiments for the 5-min (5 µL/min) and 11.5-min (1.5 µL/min) gradient used 60 variable windows (Supplementary Table 2) spanning the TOF MS mass range 400-900 Da and MS/MS mass range 140-1750 Da, with Zeno trap pulsing turned on, with MS/MS accumulation times of 13 msec. Before each Zeno SWATH DIA MS cycle an additional MS1 survey scan from 400-1500 Da was recorded for 50 ms.

The 15-min (5 µL/min)-Zeno SWATH DIA experiments used 65 variable windows (Supplementary Table 3) spanning the TOF MS mass range 400-900 Da and MS/MS mass range 140-1750 Da, with Zeno trap pulsing turned on, with MS/MS accumulation times of 13 msec. Before each Zeno SWATH DIA MS cycle an additional MS1 survey scan from 400-1500 Da was recorded for 50 ms.

The 15- and 30-min (0.250 µL/min)-Zeno SWATH DIA experiments used 85 variable windows (Supplementary Table 4) spanning the TOF MS mass range 400-900 Da and MS/MS mass range 140-1750 Da, with Zeno trap pulsing turned on, with MS/MS accumulation times of 16 ms (15-min) and 18 ms (30-min). Before each Zeno SWATH DIA MS cycle an additional MS1 survey scan from 400-1500 Da was recorded for 50 ms.

#### ZT Scan DIA

For the 5-min (5 µL/min) and 11.5-min (1.5 µL/min) ZT Scan DIA experiments, 5.1 Da Q1 isolation windows were scanned at 509 Da/s, resulting in an MS2 accumulation time of 10 ms. For the 15-min (5 µL/min) ZT Scan DIA experiments, 10.3 Da Q1 isolation windows were scanned at 514 Da/s, resulting in an MS2 accumulation time of 20 ms. For both gradients, the TOF MS and MS/MS mass range was set to 400-900 Da and 140-1750 Da, respectively. For the 15- and 30-min ZT Scan DIA experiments at 0.250 µL/min, 5 and 5.3 Da Q1 isolation windows were scanned at 298 and 267 Da/s, respectively, resulting in an MS2 accumulation time of 16.8 and 19.9 ms. For both gradients, the TOF MS and MS/MS mass range was set to 400-1500 Da and 140-1750 Da, respectively.

In Sciex OS 4.0, the acquired ZT Scan DIA raw data was calibrated using residual precursor signals and converted to SCIEX’s proprietary .wiff and .wiff.scan file formats.

#### Thermo Orbitrap Astral (Nano flow LC)

#### 1) 50 ng

A Thermo Orbitrap Astral (Thermo Fisher Scientific, Waltham, Massachusetts, United States) was coupled to a Vanquish Neo LC instrument (Thermo Fisher Scientific). Peptides were loaded directly onto the analytical column and were separated by reversed-phase chromatography using a 50 cm μPAC™ column (Thermo Fischer Scientific), featuring a structured pillar array bed with a 180 µm bed width. Buffer A consisted of 0.1% FA in water, while Buffer B contained 0.1% FA in 80% acetonitrile.

##### 15-min gradient

Chromatographic gradient was initiated at 96% buffer A and 4% buffer B at a flow rate of 750 nL/min during 1 minute. The flow rate was then reduced to 250 nL/min, and the percentage of buffer B was further increased to 40% over 15 minutes.

##### 5-min gradient

Chromatographic gradient was initiated with 96% buffer A and 4% buffer B at a flow rate of 750 nL/min during 1 minute. The flow rate was then reduced to 250 nL/min, and the percentage of buffer B was further increased to 40% over 5 minutes.

##### DDA

The mass spectrometer was operated in positive ionization mode with data-dependent acquisition, with a full MS scans over a mass range of m/z 380-980 with detection in the Orbitrap at a resolution of 180,000 and a fixed cycle time of 0.5 s. Precursor ion selection width was kept at 2-Th and a normalized collision energy of 30% was used for higher-energy collisional dissociation (HCD) fragmentation. MS2 scan range was set from 145 to 1450 m/z with detection in the Astral and maximum fill time of 2.5 ms. Dynamic exclusion was enabled and set to 10 s.

##### DIA

The mass spectrometer was operated in positive ionization mode with data-independent acquisition, with a full MS scans over a mass range of m/z 380-980 with detection in the Orbitrap at a resolution of 240,000. In each cycle of data-independent acquisition, 300 windows of 2 Th were used to isolate and fragment all precursor ions from 380 to 980 m/z. A normalized collision energy of 25% was used for HCD fragmentation. MS2 scan range was set from 150 to 2000 m/z with detection in the Astral with a maximum injection time of 3 ms.

#### 2) 50 ng – refined conditions

##### Ion Optics Direct Injection

The Orbitrap Astral (Thermo Scientific) was coupled to a Vanquish Neo UHPLC instrument (Thermo Scientific). Peptides were separated by reversed-phase chromatography using an Aurora® Elite XT (15 cm x 75 µm) (IonOpticks) column to accommodate both 5 and 15mins separations. Buffer A consisted of 0.1% FA in water, while Buffer B contained 0.1% FA in 80% acetonitrile.

Peptides were separated using a 5 and 15-min linear gradient from 3-38%B at a flow rate of 0.55 and 0.45 µL/min respectively.

##### DIA

The mass spectrometer was operated in positive ionization mode with narrow-window data-independent acquisition, with a full MS scan over a mass range of 400-800 (5-min) and 400-900 m/z (15-min) in the Orbitrap, at a resolution of 180,000 (5-min)| 240,000 (15-min). For the 5-min gradient, 133 windows of 3 Th were used to isolate and fragment all precursor ions from m/z 400 to 800, while for the 15-min gradient, 166 windows of 3 Th were used to isolate and fragment all precursor ions from m/z 400 to 900. A normalized collision energy of 25% was used for HCD fragmentation. MS2 scan range was set from 150 to 1700 m/z with detection in the Astral with a maximum injection time of 5 and 3 ms for the 5- and 15-min gradient, respectively. AGC targets were set to 500% (for MS1 scans) and 100% (for MS2 scans) for both methods, the RF lens was set to 45%, and a loop control of 0.4s (for 180,000 MS1 res) or 0.6s (for 240,000 MS1 res) ruled the cycle time.

##### uPAC Neo plus

An Orbitrap Astral (Thermo Scientific) was coupled to a Vanquish Neo UHPLC (Thermo Scientific). Peptides were loaded onto a μPAC™ Neo Trapping Column (*COL-TRPNANO16G1B2*, Thermo Scientific), and separated by reversed-phase chromatography on the latest generation μPAC™ column, a 50 cm μPAC™ Neo Plus HPLC column(*COL-UPAC050NAN*, Thermo Scientific) featuring a structured pillar array bed with a 180 µm bed width. To minimize post-column dispersion, the column was hosted in an external column oven (Sonation, PRSO-V2-PF) directly mounted onto an EASY-Spray source (Thermo Scientific) and connected via a μPAC™ Neo Plus voltage spacer (ACC-VSP1006, Thermo Scientific) to an EASY-Spray™ Nano Emitter (ES993, Thermo Scientific).

Buffer A consisted of 0.1% FA in water, while Buffer B contained 0.1% FA in 80% acetonitrile.

##### 15-min gradient

The chromatographic gradient was initiated at 4% buffer B at a flow rate of 750 nL/min, ramping to 12.5% over 3.1 minutes. The flow rate was then reduced to 450 nL/min while buffer B increased to 22.5% over 6 minutes, followed by 40% over 3.75 minutes. Finally, the column was regenerated using 90%B for 5.5 minutes.

##### 5-min gradient

The chromatographic gradient was initiated at 1 to 8% buffer B in 0.1 minutes at a flow rate of 750 nL/min, ramping to 22.4% over 3.7 minutes. The flow rate was then reduced to 650 nL/min, and buffer B was further increased to 40% over 1.25 minutes. Finally, the column was regenerated using 90%B for 4 minutes.

##### DIA

The mass spectrometer was operated in positive ionization mode with narrow-window data-independent acquisition, with a full MS scan over a mass range of 400-900 in the Orbitrap, at a resolution of 180,000 (5-min) and 240,000 (15-min), respectively. For the 5-min gradient, 166 windows of 3 Th were used to isolate and fragment all precursor ions from m/z 400 to 900, while for the 15-min gradient, 250 windows of 2 Th were used to isolate and fragment all precursor ions from m/z 400 to 900. A normalized collision energy of 25% was used for HCD fragmentation. MS2 scan range was set from 150 to 1700 m/z with detection in the Astral and a maximum injection time of 5 ms (5 min) or 3.5 ms (15 min). AGC targets were set to 500% (for MS1 scans) and 100% (for MS2 scans) for both methods, the RF lens was set to 45%, and a loop control of 0.4s (for 180,000 MS1 res) or 0.6s (for 240,000 MS1 res) ruled the cycle time.

#### 3) 250 pg

An Orbitrap Astral (Thermo Scientific) was coupled to a Vanquish Neo UHPLC instrument (Thermo Scientific). Peptides were loaded onto a μPAC™ Trapping Column (Thermo Fisher Scientific) and were separated by reversed-phase chromatography using a 50 cm μPAC™ column (Thermo Scientific), featuring a structured pillar array bed with a 180 µm bed width. Buffer A consisted of 0.1% FA in water, while Buffer B contained 0.1% FA in 80% acetonitrile.

##### 15-min gradient

Chromatographic gradient was initiated at 96.3% buffer A and 3.7% buffer B at a flow rate of 600 nL/min for 1 minute. The flow rate was then reduced to 200 nL/min, and the percentage of buffer B was further increased to 37.5% over 15 minutes.

##### 5-min gradient

Chromatographic gradient was initiated with 96.3% buffer A and 3.7% buffer B at a flow rate of 600 nL/min for 1 minute. The flow rate was then reduced to 200 nL/min, and the percentage of buffer B was further increased to 37.5% over 5 minutes.

For MS measuring, the Orbitrap Astral mass spectrometer equipped with a FAIMS Pro Duo interface (Thermo Scientific) and an EASY-Spray source was coupled to the LC. A compensation voltage of –45 V was used.

##### DDA

The mass spectrometer was operated in positive ionization mode with data-dependent acquisition, with a full MS scans over a mass range of 350-1400 m/z with detection in the Orbitrap at a resolution of 240,000 and a fixed cycle time of 1 s. Precursor ion selection width was kept at 2 Th and a normalized collision energy of 30% was used for HCD fragmentation. MS2 scan range was set from 100 to 2500 m/z with detection in the Astral and a maximum fill time of 100 ms. Dynamic exclusion was enabled and set to 20 s.

##### DIA

The mass spectrometer was operated in positive ionization mode with data-independent acquisition, with a full MS scan over a mass range of m/z 400-800 with detection in the Orbitrap at a resolution of 240,000. In each cycle of data-independent acquisition, 20 windows of 20 Th were used to isolate and fragment all precursor ions from 400 to 800 m/z. A normalized collision energy of 25% was used for HCD fragmentation. MS2 scan range was set from 150 to 2000 m/z with detection in the Astral with a maximum injection time of 100 ms.

#### 4) 250 pg – refined conditions

##### Ion Optics Direct injection

The Orbitrap Astral (Thermo Scientific) was coupled to a Vanquish Neo UHPLC instrument (Thermo Scientific). Peptides were separated by reversed-phase chromatography using an Aurora® Elite XT (15 cm x 75 µm) (IonOpticks) column to accommodate both 5 and 15 mins separations. Buffer A consisted of 0.1% FA in water, while Buffer B contained 0.1% FA in 80% acetonitrile.

Peptides were separated using a 5 and 15-min linear gradient from 3-30%B at a flow rate of 0.55 or 0.3 µL/min respectively.

#### DIA

The parameters from this section were applied for both gradients. The mass spectrometer was operated in positive ionization mode using FAIMS at CV-50 and a total carrier gas flow at minimum, or 3.5L/min. The full MS scan was covered over a mass range of 400-800 in the Orbitrap, at a resolution of 240,000. Windows of 20 Th were used to isolate and fragment all precursor ions from m/z 400 to 800, and a normalized collision energy of 25% was used for HCD fragmentation. MS2 scan range was set from 150 to 1700 m/z with detection in the Astral with a maximum injection time of 40ms. AGC targets were set to 500% (for MS1 scans) and 800% (for MS2 scans), the RF lens was set to 45%, and a loop control or 0.6s ruled the cycle time.

#### Bruker TimsTOF Ultra 2

##### 1) 250 pg

A Bruker timsTOF Ultra 2 (Bruker Daltonics, Bremen, Germany) was coupled to a nanoElute 2 LC instrument (Bruker Daltonics). Peptides were separated by reversed-phase chromatography using a 5 cm Aurora Rapid CSI column (IonOpticks) featuring a 75 µm inner diameter, 1.7 µm particle size, and 120 Å pore size. Buffer A consisted of 0.1% FA in water, while Buffer B contained 0.1% FA in acetonitrile.

##### 15-min gradient

Chromatographic gradient was initiated at 97% buffer A and 3% buffer B at a flow rate of 250 nL/min for 1 minute. The percentage of buffer B was further increased to 30% over 15 minutes.

##### 5-min gradient

Chromatographic gradient was initiated at 97% buffer A and 3% buffer B at a flow rate of 250 nL/min for 1 minute. The percentage of buffer B was further increased to 30% over 5 minutes.

For MS measuring, the timsTOF Ultra mass spectrometer (Bruker Daltonics) equipped with a CaptiveSpray nanoelectrospray source was coupled to the LC. An electrospray voltage of 1.3 kV was applied for ionization.

Data-Dependent Acquisition Parallel Accumulation Serial Fragmentation (dda-PASEF)

The mass spectrometer was operated in positive ionization mode using dda-PASEF with 10 PASEF MS/MS ramps. A full MS scan over a mass range of 100-1700 m/z and an ion mobility range from 0.64 to 1.45 1/K0 (V.s/cm2) were set, resulting in a cycle time of 1.18 s. Precursor ion selection width was kept at 2-Th@700 m/z and 3-Th @800 m/z. Ion accumulation and ramp time were set to 100 ms, and the collision energy was applied as a linear ramp from 20 eV at 1/K0 = 0.60 V.s/cm2 to 59 eV at 1/K0 =

1.6 V.s/cm2. MS2 scan range was set from 100 to 1700 m/z, the target intensity value was set to 20,000 with an intensity threshold of 1500.

###### Data-Independent Acquisition Parallel Accumulation Serial Fragmentation (dia-PASEF)

The mass spectrometer was operated in positive ionization mode using dia-PASEF, with a full MS scan over a mass range of m/z 100-1700 and an ion mobility range from 0.64 to 1.45 1/K0 (V.s/cm2) resulting in a cycle time of 0.96 s. In each DIA cycle, MS2 scan windows ranged from 400 to 1000 m/z using 25 m/z isolation windows (Supplementary Table 5). Ion accumulation and ramp time were set to 100 ms, and the collision energy was applied as a linear ramp from 20 eV at 1/K0 = 0.60 V.s/cm2 to 59 eV at 1/K0 = 1.6 V.s/cm2.

###### Diagonal Parallel Accumulation Serial Fragmentation (diagonal-PASEF)

The mass spectrometer was operated in positive ionization mode using diagonal-PASEF, with a full MS scan over a mass range of 50-3000 m/z and an ion mobility range from 0.64 to 1.45 1/K0 (V.s/cm2), resulting in a cycle time of 0.749 s.

In each cycle MS2 scan range was set from a m/z start of 400 to 850 and a m/z end from 550 to 1000, and an ion mobility range from 0.84 to 1.15 1/K0 (V.s/cm2). The number of MS2 scans was set to 6 using 25 Th isolation windows. Ion accumulation and ramp time were set to 100 ms, and the collision energy was applied as a linear ramp from 20 eV at 1/K0 = 0.60 V.s/cm2 to 59 eV at 1/K0 = 1.6 V.s/cm2.

#### 2) 50 ng

A Bruker timsTOF Ultra 2 (Bruker Daltonics, Bremen, Germany) was coupled to a nanoElute LC instrument (Bruker Daltonics). Peptides were separated by reversed-phase chromatography using a 5 cm Aurora Rapid column (IonOpticks) featuring a 150 µm inner diameter, 1.7 µm particle size, and 120 Å pore size. Buffer A consisted of 0.1% FA in water, while Buffer B contained 0.1% FA in acetonitrile.

##### 15-min gradient

Chromatographic gradient was initiated at 97% buffer A and 3% buffer B at a flow rate of 200 nL/min for 1 minute. The percentage of buffer B was further increased to 30% over 15 minutes. The column was washed with 80% B for 2 min and afterwards the column was re-equilibrated to starting conditions for an additional 6 min.

##### 5-min gradient

Chromatographic gradient was initiated at 97% buffer A and 3% buffer B at a flow rate of 200 nL/min for 1 minute. The percentage of buffer B was further increased to 30% over 5 minutes. The column was washed with 80% B for 2 min and afterwards the column was re-equilibrated to starting conditions for an additional 6 min.

For MS measuring, the timsTOF Ultra 2 mass spectrometer (Bruker Daltonics) equipped with a CaptiveSpray nanoelectrospray source was coupled to the LC. An electrospray voltage of 1.6 kV was applied for ionization.

In all methods, inverse mobilities [1/K0] from 0.64 Vs/cm^2^ to 1.45 Vs/cm^2^ with an ion accumulation and ramp time of 100 ms, respectively.

##### ddaPASEF

For ddaPASEF, one survey TIMS-MS scan was followed by ten PASEF ramps for MS/MS acquisition,wherein precursors with charge state 0 to 5 were selected for fragmentation using an intensity threshold of 500 and a target intensity of 20,000, resulting in a 1.18 s cycle time. MS/MS isolation windows were set to m/z 2.0 for precursor ions below m/z 700 and m/z 3.0 for those above. MS/MS spectra were scanned from *m/z* 100 to *m/z* 1700. Active Exclusion was enabled and set to 0.40 min.

##### diaPASEF

The mass spectrometer was operated in positive ionization mode using dia-PASEF, with a full MS scan over a mass range of m/z 100-1700 and an ion mobility range from 0.64 to 1.37 1/K0 (V.s/cm2) resulting in a cycle time of 0.96 s. In each DIA cycle, MS2 scan windows ranged from 400 to 1000 m/z using 25 m/z isolation windows (Supplementary Table 5). The collision energy was applied as a linear ramp from 20 eV at 1/K0 = 0.60 V.s/cm2 to 63 eV at 1/K0 = 1.6 V.s/cm2.

##### diagonal-PASEF

The mass spectrometer was operated in positive ionization mode using diagonal-PASEF, with a full MS scan over a mass range of 100-1700 m/z and an ion mobility range from 0.64 to 1.45 1/K0 (V.s/cm2), resulting in a cycle time of 0.64 s.

In each cycle MS2 scan range was set from a m/z start of 400 to 980 and a m/z end from 550 to 1130, and an ion mobility range from 0.82 to 1.25 1/K0 (V.s/cm2). The number of MS2 scans was set to 6 using 30 Th isolation windows. The collision energy was applied as a linear ramp from 20 eV at 1/K0 = 0.60 V.s/cm2 to 59 eV at 1/K0 = 1.6 V.s/cm2.

### Data Records

Data record 1. The mass spectrometry proteomics data have been deposited to the ProteomeXchange Consortium (http://proteomecentral.proteomexchange.org) via the PRIDE partner repository ^39^ with the dataset identifiers, PXD070049 ^40^ and PXD071205 ^41^. A Sample and Data Relationship File (SDRF) and an Investigation Description File (IDF) have been uploaded to ProteomeXchange ^42,43^. These files are used to annotate the sample metadata and link the metadata to the corresponding data file(s) and thus will improve the reproducibility and reanalysis of this comprehensive benchmark dataset. Log in to the PRIDE website using the following project accession: PXD070049 (token: hcXc5dENPPrf). Alternatively, reviewer(s) can access the dataset by logging in to the PRIDE website using the following username: (Password: LF0Bpyt0gt1g). The refined Astral datasets can be accessed through PRIDE using the following project accession: PXD071205 (token: 5Er73LxcO8VH). Alternatively, reviewer(s) can access the dataset by logging in to the PRIDE website using the following username: (Password: KXlk44y48XYF)

Data record 2. The AutoQC data analysed in Skyline is available from Panorama Public with the link https://panoramaweb.org/LFQBenchmark_Generation_Beta.url ^44^.

### Technical Validation

Similar to the first generation LFQ benchmark dataset, a continuous system suitability monitoring protocol was implemented to assess LC-MS/MS performance over time. A commercial *E. coli* protein digest extract was used as automatic quality control (AutoQC) sample across most of the instruments. These samples were acquired in DDA mode using a 15-minute LC gradient, with the exception of the 50 ng timsTOF Ultra 2, where 5- and 15-minute gradients were alternated intermittently.

System suitability was evaluated using peptide-identification-free metrics—such as retention time, peak area, and mass accuracy—extracted via the vendor-neutral Panorama AutoQC framework ^45^. To define suitable peptides, a subset of *E. coli* samples per instrument was peak picked using MSConvert (v3.0.23229) and searched with Mascot Daemon (v2.8.0) against an *E. coli* FASTA database using the following parameters: (i) 20 ppm precursor mass tolerance, (ii) 50 ppm fragment mass tolerance, and (iii) up to two missed cleavages.

Mascot identification results were exported as .dat files and imported into Skyline Daily (v24.1). Four highly abundant proteins—P0A6F3, P0A6F5, P0A6Y8, and P0A853—were retained. Low-intensity and co-eluting precursors were excluded. AutoQC Loader (v24.1.0.443) was configured per instrument to automatically import raw files containing “AutoQC” in the filename pattern. Processed results and reports are available in the PanoramaWeb folder: U of Ghent Pharma Biotech Lab – LFQBenchmark_Generation_Beta.

Notably, far fewer outliers were observed compared to the previous generation benchmark dataset when evaluating TIC and precursor area, retention time, and mass accuracy (Supplementary Figure 1). The improvement in performance may be attributed to the use of recent workflows, much shorter LC gradients, as well as the enhanced robustness of the newer generation LC and MS instruments. As inherent variations in chromatographic setups would directly influence identifications, raw files were also visually inspected to ensure data quality. A set of chromatograms representative of the pool of separations is available in Supplementary Figure 2 and 3.

For the timsTOF Ultra 2 – 250 pg dataset, a decrease in precursor area was seen over the time course of the sample batch. Whether the observed effect stems from a true loss in instrument sensitivity or from sample-related issues cannot be definitively confirmed. However, since the overall TIC area remained stable, instrument degradation is likely not the primary cause.

The 250 pg samples on the Orbitrap Astral were measured using a stock concentration of 5 ng/µL, with injection volumes of 0.05 µL, which is well established and supported by the Vanquish Neo LC system. Nonetheless, a general decrease in instrument sensitivity was observed on the Orbitrap Astral system used for the 250 pg batch, as evidenced by lower precursor intensities and reduced total ion chromatogram (TIC) areas from overlapped chromatograms, compared to earlier runs. This could suggest a partial workflow performance decline during the analysis, or more likely, an effect caused by the ultra-low load nature of the acquisitions which may have contributed to the reduced identification depth in this specific dataset.

For the ZenoTOF 8600 (5 µL/min) and Orbitrap Astral 50 ng (refined conditions) dataset, a different quality control procedure was applied. More specifically, for the ZenoTOF 8600 the *E. coli* protein digest was measured in the respective LC gradient and DIA methodology employed during the acquisition of the different sample batches. Therefore, we were no longer able to use Mascot as search engine and opted for DIA-NN (v2.1.0) for identification purposes. As expected, the reproducibility in number of precursor and protein group identifications within batch was extremely high which corroborates with the AutoQC DDA results obtained for the ZenoTOF 7600+. For the refined 50 ng Orbitrap Astral dataset, we selected a more complex proteome, namely a HeLa protein digest standard (Thermo Fischer Scientific, Waltham, Massachusetts, United States), and analyzed it using DIA to expand the number of target peptides available for QC monitoring and again achieved extremely high reproducibility, with 98.5% data completeness when performing directDIA+ analysis in Spectronaut.

In summary, the implemented system suitability monitoring confirmed that the dataset is of high quality and representative of realistic performance achieved in modern proteomics core facilities.

### Usage Notes

In the following section, we illustrate the potential of this curated dataset through selected use cases, without positioning it as a benchmarking exercise of instrument performance. The core dataset was generated using unified operating conditions, rather than instrument-specific performance tuning. Although additional analyses were performed under refined configurations, these serve as complementary examples. Therefore, the dataset should not be used to draw conclusions about which instrument performs “best.” Instead, it enables controlled, transparent, and methodologically fair evaluations of tools, workflows, and acquisition strategies.

A central use case for this dataset is its integration into ProteoBench, a community-driven benchmarking platform that inspired the development of this second-generation LFQ reference dataset. ProteoBench provides a transparent, reproducible environment for evaluating proteomics data analysis software. It also aims to facilitate open discussions on evaluation principles, for example on how benchmarking datasets, should be composed. ProteoBench enables users and software developers to perform targeted, side-by-side comparisons ^38^. The diversity of instruments and acquisition strategies represented in this dataset makes it well-suited as a reasonable resource for assessing identification overlap, quantification error, and workflow robustness across proteomics data analysis tools. For example, precursor and protein identifications and their quantification accuracy can be compared across DIA pipelines such as AlphaDIA ^46^, DIA-NN ^47^, FragPipe ^48^, PEAKS ^49^ and Spectronaut (Supplementary Figure 4 and 5).

Since this dataset includes data acquired across multiple MS instruments, LC setups and acquisition methodologies, it also provides a unique resource to evaluate how technological innovations and parameter tweaking affect proteomics performance. Firstly, the use of a scanning quadrupole for DIA has gained momentum across multiple vendors: SCIEX introduced ZT Scan DIA^27^ with the ZenoTOF 7600+, while Bruker implemented a similar concept into the timsTOF platform as miDIA-PASEF^29^ and synchro-PASEF^30^ (collectively referred to as diagonal-PASEF^28^). These approaches have been shown to improve precursor selectivity by incorporating movement of the Q1 isolation window over time, effectively adding an additional separation dimension to DIA acquisition. The advantage of these approaches have been especially important in short-gradient, low-input experiments ^28^. Here, we evaluated performance trends previously reported in the literature and found them reflected in our dataset. In the low-input 250 pg Ultra 2 dataset, we observe a clear improvement in quantification accuracy when using diagonal-PASEF compared to classical dia-PASEF (Supplementary Figure 6), reflected in the lower median deviation from the expected log_2_FCs. This effect is consistent across both the 5- and 15-minute gradients, although it is accompanied by a reduction in the number of quantified precursors. It remains unclear whether this reduction stems from suboptimal DIA-NN parameter settings or if it is consistent accross the latest generation of TIMS instrument, as we were unable to evaluate a 250 pg dataset on the timsUltra AIP in this study. Similarly, for the ZenoTOF 8600 datasets, ZT Scan DIA acquisition improves quantification relative to Zeno SWATH DIA for the 5- and 11-minute gradients using microflow LC, whereas this benefit is not observed for the longer 15-minute gradient (Supplementary Figure 7). Using nanoflow LC, on the other hand, ZT Scan DIA acquisition provides better quantification for 15-minute gradients as well.

Secondly, and also shown by this last observation, this dataset allows for the investigation of how chromatographic setups impact identification performance. For many years, SCIEX users have favored microflow LC (∼5 µL/min) as a robust and reproducible alternative to nanoflow separations, despite its known sensitivity trade-offs ^50^. Given the rising importance of sensitivity in low-input samples, especially single-cell proteomics, we systematically compared three flow rates—5.0, 1.5, and 0.25 µL/min—on the ZenoTOF 8600 at 50 ng (Supplementary Figure 8). Identification rates scaled inversely with flow rate, when moving from microflow to low-nanoflow, with intermediate results at 1.5 µL/min. Gains were most pronounced at the precursor level, consistent with improved ionization efficiency and reduced ion suppression at lower flow rates ^51^. At the protein group level, increases were more modest, reflecting redundancy in peptide-to-protein mapping.

It should be emphasized that the main aim of this dataset is not to maximize performance on each MS platform, but rather to generate a representative dataset that reflects average instrument usage conditions. Nonetheless, recognizing that acquisition parameters can be further optimized and that doing so is expected to enhance performance, we acquired a set of two additional 50 ng and one 250 pg datasets on the Orbitrap Astral under more refined analytical conditions.

Across the two refined 50 ng Orbitrap Astral datasets, we observed clear performance improvements compared to the standardized acquisitions (Supplementary Figure 9). Notably, the nature of these improvements differed depending on the refinement strategy. In one of these, the most vital change concerned the standardized linear LC gradient from 3–30%, which was optimized to achieve a more uniform distribution of peptide separation across the chromatographic run. This workflow was also operated in a trap-and-elute setup that minimizes post-column dispersion, a feature enabled by the latest generation μPAC Neo Plus column, which is particularly advantageous for continuous measurements and for meeting laboratory robustness requirements. For the 15-min gradient, this configuration substantially increased the number of quantified precursors, albeit at the cost of higher quantification error. By contrast, a refined direct-injection configuration yielded a more moderate gain in sensitivity but did not introduce additional quantification error. For the 5-minute gradient, both refined methods improved quantification accuracy, while the increase in quantified precursors was more modest, likely limited by the narrower m/z scan range (400-800 vs. 380-980). A MS2 quantitative strategy was opted for this study, and further quantitative improvements might have been observed on the short gradient while using an MS1 strategy. Importantly, the improvements presented here are not exhaustive. Further optimization, such as fine-tuning LC gradients and workflow, or operating FAIMS at lower resolution for low-input samples may provide additional gains in sensitivity, sampling efficiency, and quantitative performance ^52^. Collectively, these results highlight that substantial performance benefits can be unlocked when LC-MS parameters are carefully tailored to the analytical objectives and sample amount.

It should be noted that all LC-MS platforms included in this study are capable of such improved performance through similar fine-tuning. Again, the data presented here were intentionally generated under standardized instructions and to the best efforts of the operators. Consequently, no direct conclusions can be drawn regarding which platform is “best” in terms of identification depth or quantitative performance.

By spanning different instruments, acquisition methods, and LC gradients, this dataset provides a controlled yet representative resource for fair and transparent evaluations across tools and platforms. This enables assessment of performance metrics, whether in the context of software benchmarking through initiatives such as ProteoBench, the evaluation of novel acquisition approaches like narrow-window DIA, diagonal-PASEF and ZT Scan DIA, or the study of methodological variables such as workflows, flow rate and MS parameter optimization. In the long term, such community-driven benchmarking efforts will not only drive continuous improvements in both bioinformatics pipelines and experimental practices but also empower end-users, including biologists, to select the strategies and tools that best match their research needs and expectations.

## Code Availability

The Jupyter notebooks used to generate Figure 2 and the Supplementary Figures 4, 5, 6, 7, 8 and 9 are available through Zenodo, under 10.5281/zenodo.17936656 (zenodo.org/records/17936657).

## Acknowledgements

This research was funded by grants from the Research Foundation Flanders (FWO) and the Special University Research Fund of Ghent University awarded to B.VP (grant numbers: 1278023N and BOF/PDO/2025/049). R.D., L.M. and R.B. acknowledge funding from the Research Foundation Flanders (FWO) [1SH9O24N, G010023N, 12A6L24N]. L.M. acknowledges funding from the Horizon Europe Projects BAXERNA 2.0 [101080544] and COMBINE [101191739], and from the Ghent University Concerted Research Action [BOF21/GOA/033]. L.M. and C.Car are further supported by the CHIST-ERA project ODEEP-EU [G0GDV23N]. We thank ProGenTomics for providing access to the ZenoTOF 7600+ system used in this study. YPR thanks European Molecular Biology Laboratory core funding, Wellcome grants (208391/Z/17/Z and 223745/Z/21/Z) and the BBSRC grant ‘DIA-Exchange’ (BB/X001911/1). This work was supported by the Agence Nationale de la Recherche via the French Proteomic Infrastructure (ProFI UAR2048; ANR-10-INBS08-03 & ANR-24-INBS-0015), by the Region Grand-Est (SC-Proteomics project) and by the ITMO Cancer of Aviesan within the framework of the 2021-2030 Cancer Control Strategy, on funds administered by Inserm (ProteomiSC project), for equipment funds at C.Car. C. Car and C.K. acknowledge the Interdisciplinary Thematic Institute IMS, the drug discovery and development institute, as part of the ITI 2021-2028 program of the University of Strasbourg, CNRS and Inserm, supported by IdEx Unistra (ANR-10-IDEX-0002), and by SFRI-STRAT’US project (ANR-20-SFRI-0012). We would like to thank BSI for their support in setting up the analysis in PEAKS.

## Author contributions

B.VP, R.D., R.B., and M.D. conceived the study and wrote the draft manuscript with contributions from all authors. B.VP prepared the samples and performed the ZenoTOF 7600+ data acquisition with the help from I.B. and JB.V.; P.P. performed the ZenoTOF 8600 data acquisition; C.C., E.S., J.B.R., C.K., C.Car, F.F., L.L., P.M. and B.G. performed the Orbitrap Astral data acquisition; J.B.R., C.K., C.Car performed the timsTOF Ultra 2 – 250 pg data acquisition; S.P., C.P. and P.N. performed the timsTOF Ultra 2 – 50 ng data acquisition and R.D. and R.B. wrote the custom scripts for data analysis. YPR organised the ProteomeXchange submission. DD and LM co-supervised the experiment.

