## Supplementary Information for "LFQ Benchmark Dataset - Generation Beta: Assessing Modern Proteomics Instruments and Acquisition Workflows with High-Throughput LC Gradients"

**Supplementary Table 1.** Standardized sample injection order scheme provided to all laboratories.

| Individual Standards |
|---|
| LFQ_NameInstrument_Acquisitionmethodology_5/15min_250pg/50ng_Standard_AutoQC_01 |
| LFQ_NameInstrument_Acquisitionmethodology_5/15min_250pg/50ng_Ecoli_01 |
| LFQ_NameInstrument_Acquisitionmethodology_5/15min_250pg/50ng_Human_01 |
| LFQ_NameInstrument_Acquisitionmethodology_5/15min_250pg/50ng_Yeast_01 |
| LFQ_NameInstrument_Acquisitionmethodology_5/15min_250pg/50ng_Standard_AutoQC_02 |
| LFQ_NameInstrument_Acquisitionmethodology_5/15min_250pg/50ng_Ecoli_02 |
| LFQ_NameInstrument_Acquisitionmethodology_5/15min_250pg/50ng_Human_02 |
| LFQ_NameInstrument_Acquisitionmethodology_5/15min_250pg/50ng_Yeast_02 |
| LFQ_NameInstrument_Acquisitionmethodology_5/15min_250pg/50ng_Standard_AutoQC_03 |
| LFQ_NameInstrument_Acquisitionmethodology_5/15min_250pg/50ng_Ecoli_03 |
| LFQ_NameInstrument_Acquisitionmethodology_5/15min_250pg/50ng_Human_03 |
| LFQ_NameInstrument_Acquisitionmethodology_5/15min_250pg/50ng_Yeast_03 |
| LFQ_NameInstrument_Acquisitionmethodology_5/15min_250pg/50ng_Standard_AutoQC_04 |

| HYE samples |
|---|
| LFQ_NameInstrument_Acquisitionmethodology_5/15min_AutoQC_01 |
| LFQ_NameInstrument_Acquisitionmethodology_5/15min_250pg/50ng_Condition_A_REP1 |
| LFQ_NameInstrument_Acquisitionmethodology_5/15min_250pg/50ng_Condition_B_REP1 |
| LFQ_NameInstrument_Acquisitionmethodology_5/15min_250pg/50ng_Condition_C_REP1 |
| LFQ_NameInstrument_Acquisitionmethodology_5/15min_250pg/50ng_Condition_A_REP2 |
| LFQ_NameInstrument_Acquisitionmethodology_5/15min_250pg/50ng_Condition_B_REP2 |
| LFQ_NameInstrument_Acquisitionmethodology_5/15min_250pg/50ng_Condition_C_REP2 |
| LFQ_NameInstrument_Acquisitionmethodology_5/15min_AutoQC_02 |
| LFQ_NameInstrument_Acquisitionmethodology_5/15min_250pg/50ng_Condition_A_REP3 |
| LFQ_NameInstrument_Acquisitionmethodology_5/15min_250pg/50ng_Condition_B_REP3 |
| LFQ_NameInstrument_Acquisitionmethodology_5/15min_250pg/50ng_Condition_C_REP3 |
| LFQ_NameInstrument_Acquisitionmethodology_5/15min_250pg/50ng_Condition_A_REP4 |
| LFQ_NameInstrument_Acquisitionmethodology_5/15min_250pg/50ng_Condition_B_REP4 |
| LFQ_NameInstrument_Acquisitionmethodology_5/15min_250pg/50ng_Condition_C_REP4 |
| LFQ_NameInstrument_Acquisitionmethodology_5/15min_AutoQC_03 |
| LFQ_NameInstrument_Acquisitionmethodology_5/15min_250pg/50ng_Condition_A_REP5 |
| LFQ_NameInstrument_Acquisitionmethodology_5/15min_250pg/50ng_Condition_B_REP5 |
| LFQ_NameInstrument_Acquisitionmethodology_5/15min_250pg/50ng_Condition_C_REP5 |
| LFQ_NameInstrument_Acquisitionmethodology_5/15min_250pg/50ng_Condition_A_REP6 |

| |
|---|
| LFQ_NameInstrument_Acquisitionmethodology_5/15min_250pg/50ng_Condition_B_REP6 |
| LFQ_NameInstrument_Acquisitionmethodology_5/15min_250pg/50ng_Condition_C_REP6 |
| LFQ_NameInstrument_Acquisitionmethodology_5/15min_AutoQC_04 |

**Supplementary Table 2.** 60 Variable Window Scheme applied for the 5min- Zeno SWATH data acquired on the SCIEX ZenoTOF 7600+.

| Start | End | CES | | Start | End | CES |
|---|---|---|---|---|---|---|
| 399.5 | 409.5 | 0 | | 736.5 | 745.5 | 0 |
| 408.5 | 418.5 | 0 | | 744.5 | 753.5 | 0 |
| 417.5 | 427.5 | 0 | | 752.5 | 761.5 | 0 |
| 426.5 | 436.5 | 0 | | 760.5 | 769.5 | 0 |
| 435.5 | 445.5 | 0 | | 768.5 | 777.5 | 0 |
| 444.5 | 454.5 | 0 | | 776.5 | 785.5 | 0 |
| 453.5 | 463.5 | 0 | | 784.5 | 793.5 | 0 |
| 462.5 | 472.5 | 0 | | 792.5 | 801.5 | 0 |
| 471.5 | 481.5 | 0 | | 800.5 | 810.5 | 0 |
| 480.5 | 489.5 | 0 | | 809.5 | 819.5 | 0 |
| 488.5 | 497.5 | 0 | | 818.5 | 828.5 | 0 |
| 496.5 | 505.5 | 0 | | 827.5 | 837.5 | 0 |
| 504.5 | 513.5 | 0 | | 836.5 | 846.5 | 0 |
| 512.5 | 521.5 | 0 | | 845.5 | 855.5 | 0 |
| 520.5 | 529.5 | 0 | | 854.5 | 864.5 | 0 |
| 528.5 | 537.5 | 0 | | 863.5 | 873.5 | 0 |
| 536.5 | 545.5 | 0 | | 872.5 | 882.5 | 0 |
| 544.5 | 553.5 | 0 | | 881.5 | 891.5 | 0 |
| 552.5 | 561.5 | 0 | | 890.5 | 900.5 | 0 |
| 560.5 | 569.5 | 0 | | | | |
| 568.5 | 577.5 | 0 | | | | |
| 576.5 | 585.5 | 0 | | | | |
| 584.5 | 593.5 | 0 | | | | |
| 592.5 | 601.5 | 0 | | | | |
| 600.5 | 609.5 | 0 | | | | |
| 608.5 | 617.5 | 0 | | | | |
| 616.5 | 625.5 | 0 | | | | |
| 624.5 | 633.5 | 0 | | | | |
| 632.5 | 641.5 | 0 | | | | |
| 640.5 | 649.5 | 0 | | | | |
| 648.5 | 657.5 | 0 | | | | |
| 656.5 | 665.5 | 0 | | | | |
| 664.5 | 673.5 | 0 | | | | |
| 672.5 | 681.5 | 0 | | | | |
| 680.5 | 689.5 | 0 | | | | |
| 688.5 | 697.5 | 0 | | | | |

| | | |
|---|---|---|
| 696.5 | 705.5 | 0 |
| 704.5 | 713.5 | 0 |
| 712.5 | 721.5 | 0 |
| 720.5 | 729.5 | 0 |
| 728.5 | 737.5 | 0 |

**Supplementary Table 3.** 65 Variable Window Scheme applied for the Zeno SWATH data acquired on the SCIEX ZenoTOF 7600+ and ZenoTOF 8600.

| Start | End | CES | | Start | End | CES |
|---|---|---|---|---|---|---|
| 399.5 | 409.5 | 0 | | 696.5 | 705.5 | 0 |
| 408.5 | 418.5 | 0 | | 704.5 | 713.5 | 0 |
| 417.5 | 427.5 | 0 | | 712.5 | 721.5 | 0 |
| 426.5 | 436.5 | 0 | | 720.5 | 729.5 | 0 |
| 435.5 | 445.5 | 0 | | 728.5 | 737.5 | 0 |
| 444.5 | 454.5 | 0 | | 736.5 | 745.5 | 0 |
| 453.5 | 463.5 | 0 | | 744.5 | 753.5 | 0 |
| 462.5 | 472.5 | 0 | | 752.5 | 761.5 | 0 |
| 471.5 | 481.5 | 0 | | 760.5 | 769.5 | 0 |
| 480.5 | 489.5 | 0 | | 768.5 | 777.5 | 0 |
| 488.5 | 497.5 | 0 | | 776.5 | 785.5 | 0 |
| 496.5 | 505.5 | 0 | | 784.5 | 793.5 | 0 |
| 504.5 | 513.5 | 0 | | 792.5 | 801.5 | 0 |
| 512.5 | 521 | 0 | | 800.5 | 810.5 | 0 |
| 520 | 528.5 | 0 | | 809.5 | 819.5 | 0 |
| 527.5 | 536 | 0 | | 818.5 | 828.5 | 0 |
| 535 | 543.5 | 0 | | 827.5 | 837.5 | 0 |
| 542.5 | 550.5 | 0 | | 836.5 | 846.5 | 0 |
| 549.5 | 557.5 | 0 | | 845.5 | 855.5 | 0 |
| 556.5 | 564.5 | 0 | | 854.5 | 864.5 | 0 |
| 563.5 | 571 | 0 | | 863.5 | 873.5 | 0 |
| 570 | 577.5 | 0 | | 872.5 | 882.5 | 0 |
| 576.5 | 583.5 | 0 | | 881.5 | 891.5 | 0 |
| 582.5 | 590 | 0 | | 890.5 | 900.5 | 0 |
| 589 | 596.5 | 0 | | | | |
| 595.5 | 601.5 | 0 | | | | |
| 600.5 | 605.5 | 0 | | | | |
| 604.5 | 609.5 | 0 | | | | |
| 608.5 | 613.5 | 0 | | | | |
| 612.5 | 617.5 | 0 | | | | |
| 616.5 | 622.5 | 0 | | | | |
| 621.5 | 628.5 | 0 | | | | |
| 627.5 | 634.5 | 0 | | | | |
| 633.5 | 640.5 | 0 | | | | |
| 639.5 | 649 | 0 | | | | |
| 648 | 657.5 | 0 | | | | |
| 656.5 | 665.5 | 0 | | | | |
| 664.5 | 673.5 | 0 | | | | |
| 672.5 | 681.5 | 0 | | | | |
| 680.5 | 689.5 | 0 | | | | |
| 688.5 | 697.5 | 0 | | | | |

Supplementary Table 4. 85 Variable Window Scheme applied for the Zeno SWATH data acquired on the SCIEX ZenoTOF 8600.

| Start | End | CES | | Start | End | CES | | Start | End | CES |
|---|---|---|---|---|---|---|---|---|---|---|
| 399.5 | 406.5 | 0 | | 613.5 | 619.5 | 0 | | 875.5 | 885.5 | 0 |
| 405.5 | 412.5 | 0 | | 618.5 | 624.5 | 0 | | 884.5 | 894.5 | 0 |
| 411.5 | 418.5 | 0 | | 623.5 | 629.5 | 0 | | 893.5 | 903.5 | 0 |
| 417.5 | 424.5 | 0 | | 628.5 | 634.5 | 0 | | | | |
| 423.5 | 430.5 | 0 | | 633.5 | 639.5 | 0 | | | | |
| 429.5 | 436.5 | 0 | | 638.5 | 644.5 | 0 | | | | |
| 435.5 | 442.5 | 0 | | 643.5 | 649.5 | 0 | | | | |
| 441.5 | 448.5 | 0 | | 648.5 | 654.5 | 0 | | | | |
| 447.5 | 454.5 | 0 | | 653.5 | 660.5 | 0 | | | | |
| 453.5 | 459.5 | 0 | | 659.5 | 666.5 | 0 | | | | |
| 458.5 | 464.5 | 0 | | 665.5 | 672.5 | 0 | | | | |
| 463.5 | 469.5 | 0 | | 671.5 | 678.5 | 0 | | | | |
| 468.5 | 474.5 | 0 | | 677.5 | 684.5 | 0 | | | | |
| 473.5 | 479.5 | 0 | | 683.5 | 690.5 | 0 | | | | |
| 478.5 | 484.5 | 0 | | 689.5 | 696.5 | 0 | | | | |
| 483.5 | 489.5 | 0 | | 695.5 | 702.5 | 0 | | | | |
| 488.5 | 494.5 | 0 | | 701.5 | 708.5 | 0 | | | | |
| 493.5 | 499.5 | 0 | | 707.5 | 714.5 | 0 | | | | |
| 498.5 | 504.5 | 0 | | 713.5 | 720.5 | 0 | | | | |
| 503.5 | 509.5 | 0 | | 719.5 | 726.5 | 0 | | | | |
| 508.5 | 514.5 | 0 | | 725.5 | 732.5 | 0 | | | | |
| 513.5 | 519.5 | 0 | | 731.5 | 738.5 | 0 | | | | |
| 518.5 | 524.5 | 0 | | 737.5 | 744.5 | 0 | | | | |
| 523.5 | 529.5 | 0 | | 743.5 | 750.5 | 0 | | | | |
| 528.5 | 534.5 | 0 | | 749.5 | 756.5 | 0 | | | | |
| 533.5 | 539.5 | 0 | | 755.5 | 763.5 | 0 | | | | |
| 538.5 | 544.5 | 0 | | 762.5 | 770.5 | 0 | | | | |
| 543.5 | 549.5 | 0 | | 769.5 | 777.5 | 0 | | | | |
| 548.5 | 554.5 | 0 | | 776.5 | 784.5 | 0 | | | | |
| 553.5 | 559.5 | 0 | | 783.5 | 791.5 | 0 | | | | |
| 558.5 | 564.5 | 0 | | 790.5 | 798.5 | 0 | | | | |
| 563.5 | 569.5 | 0 | | 797.5 | 805.5 | 0 | | | | |
| 568.5 | 574.5 | 0 | | 804.5 | 812.5 | 0 | | | | |
| 573.5 | 579.5 | 0 | | 811.5 | 819.5 | 0 | | | | |
| 578.5 | 584.5 | 0 | | 818.5 | 826.5 | 0 | | | | |
| 583.5 | 589.5 | 0 | | 825.5 | 834.5 | 0 | | | | |
| 588.5 | 594.5 | 0 | | 833.5 | 842.5 | 0 | | | | |
| 593.5 | 599.5 | 0 | | 841.5 | 850.5 | 0 | | | | |
| 598.5 | 604.5 | 0 | | 849.5 | 858.5 | 0 | | | | |
| 603.5 | 609.5 | 0 | | 857.5 | 867.5 | 0 | | | | |
| 608.5 | 614.5 | 0 | | 866.5 | 876.5 | 0 | | | | |

Supplementary Table 5. diaPASEF window scheme

| Cycle Id | Start IM [1/K0] | End IM [1/K0] | Start Mass [m/z] | End Mass [m/z] |
|---|---|---|---|---|
| **1** | 0.64 | 0.83 | 400 | 425 |
| **1** | 0.83 | 1.01 | 600 | 625 |
| **1** | 1.01 | 1.37 | 800 | 825 |
| **2** | 0.64 | 0.85 | 425 | 450 |
| **2** | 0.85 | 1.04 | 625 | 650 |
| **2** | 1.04 | 1.37 | 825 | 850 |
| **3** | 0.64 | 0.87 | 450 | 475 |
| **3** | 0.87 | 1.06 | 650 | 675 |
| **3** | 1.06 | 1.37 | 850 | 875 |
| **4** | 0.64 | 0.9 | 475 | 500 |
| **4** | 0.9 | 1.09 | 675 | 700 |
| **4** | 1.09 | 1.37 | 875 | 900 |
| **5** | 0.64 | 0.92 | 500 | 525 |
| **5** | 0.92 | 1.11 | 700 | 725 |
| **5** | 1.11 | 1.37 | 900 | 925 |
| **6** | 0.64 | 0.94 | 525 | 550 |
| **6** | 0.94 | 1.13 | 725 | 750 |
| **6** | 1.13 | 1.37 | 925 | 950 |
| **7** | 0.64 | 0.97 | 550 | 575 |
| **7** | 0.97 | 1.16 | 750 | 775 |
| **7** | 1.16 | 1.37 | 950 | 975 |
| **8** | 0.64 | 0.99 | 575 | 600 |
| **8** | 0.99 | 1.18 | 775 | 800 |
| **8** | 1.18 | 1.37 | 975 | 1000 |

a) <u>Orbitrap Astral – 50 ng</u>

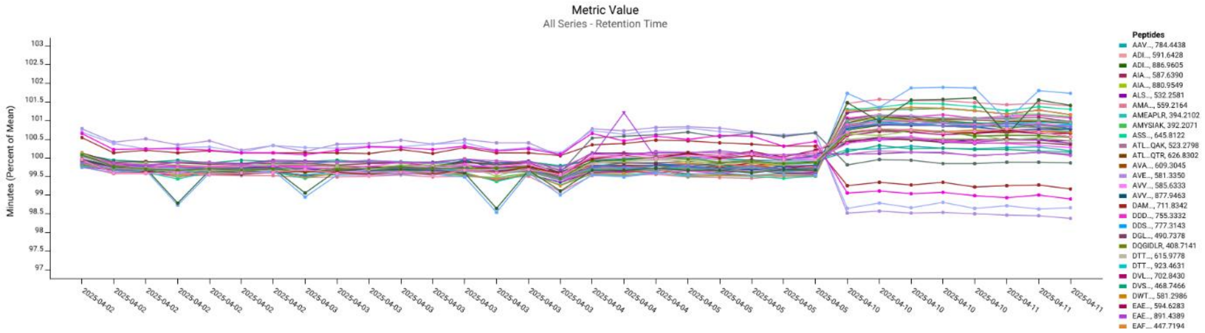

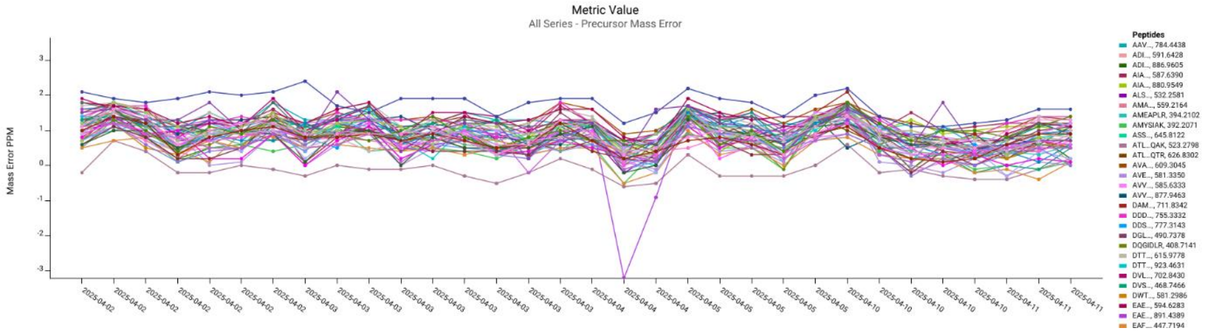

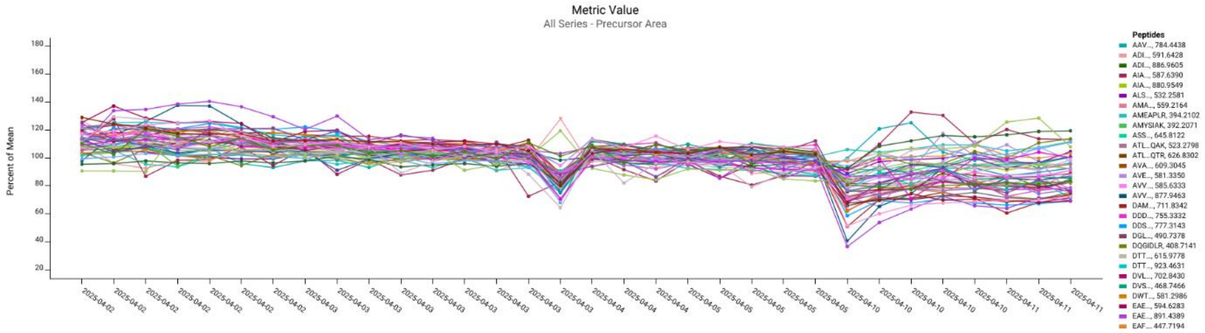

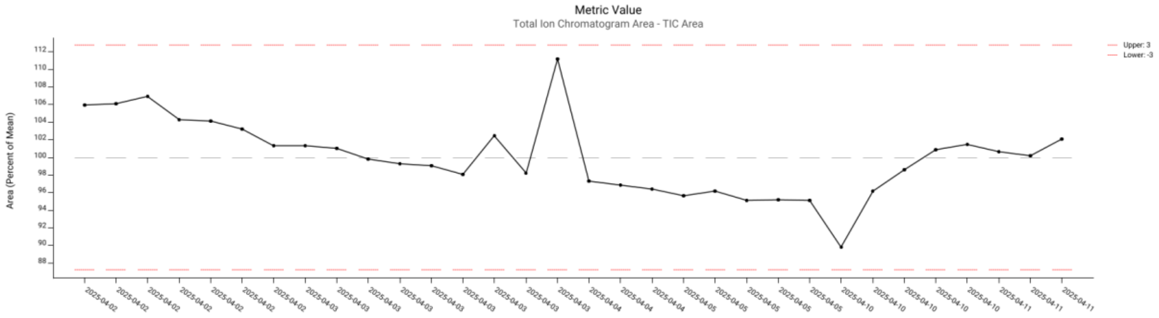

b) ZenoTOF 8600

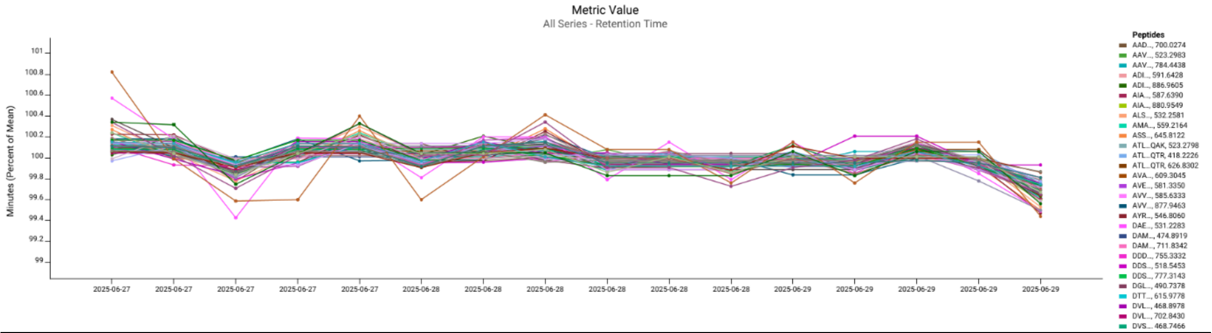

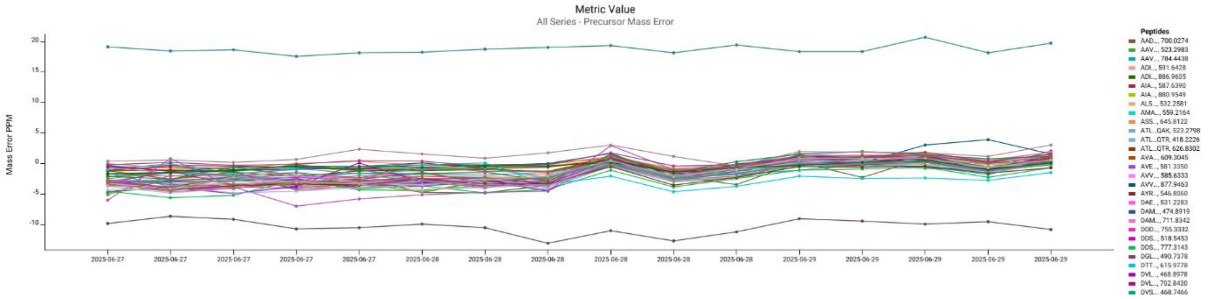

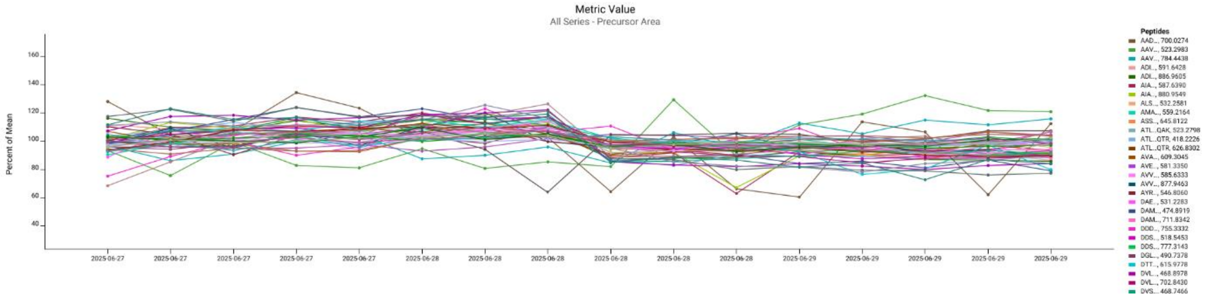

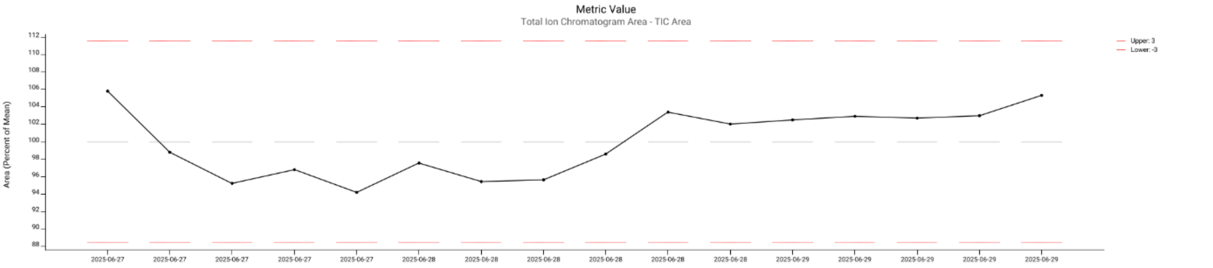

c) <u>Ultra 2</u>

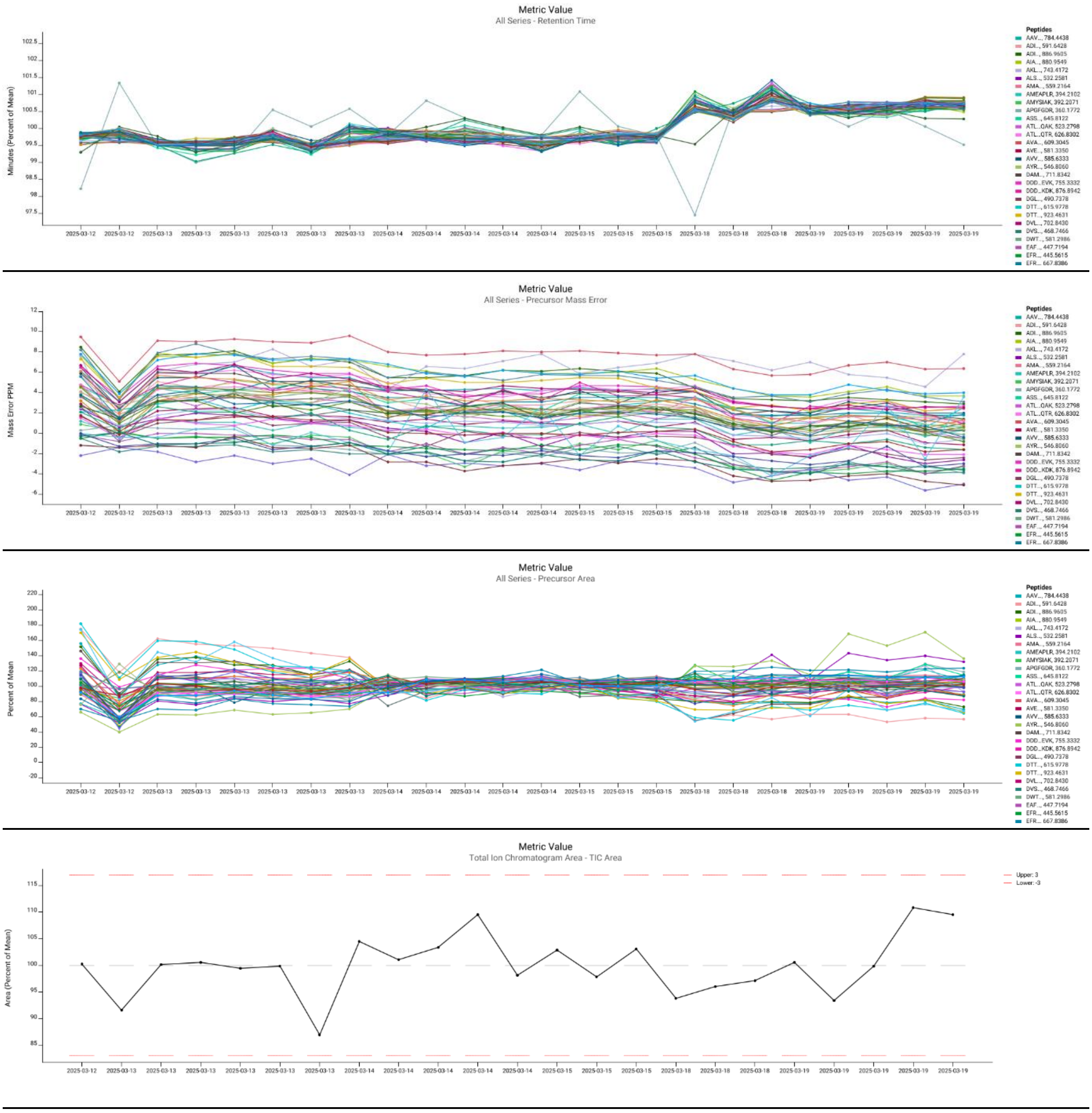

**Supplementary Figure 1.** Overview of retention time, precursor mass error, precursor area, and TIC area variability over time for the 50 ng a) Orbitrap Astral (standard conditions) b) ZenoTOF 8600 (microflow) and c) timsTOF Ultra 2 dataset. Figures were generated using Skyline Panorama QC.

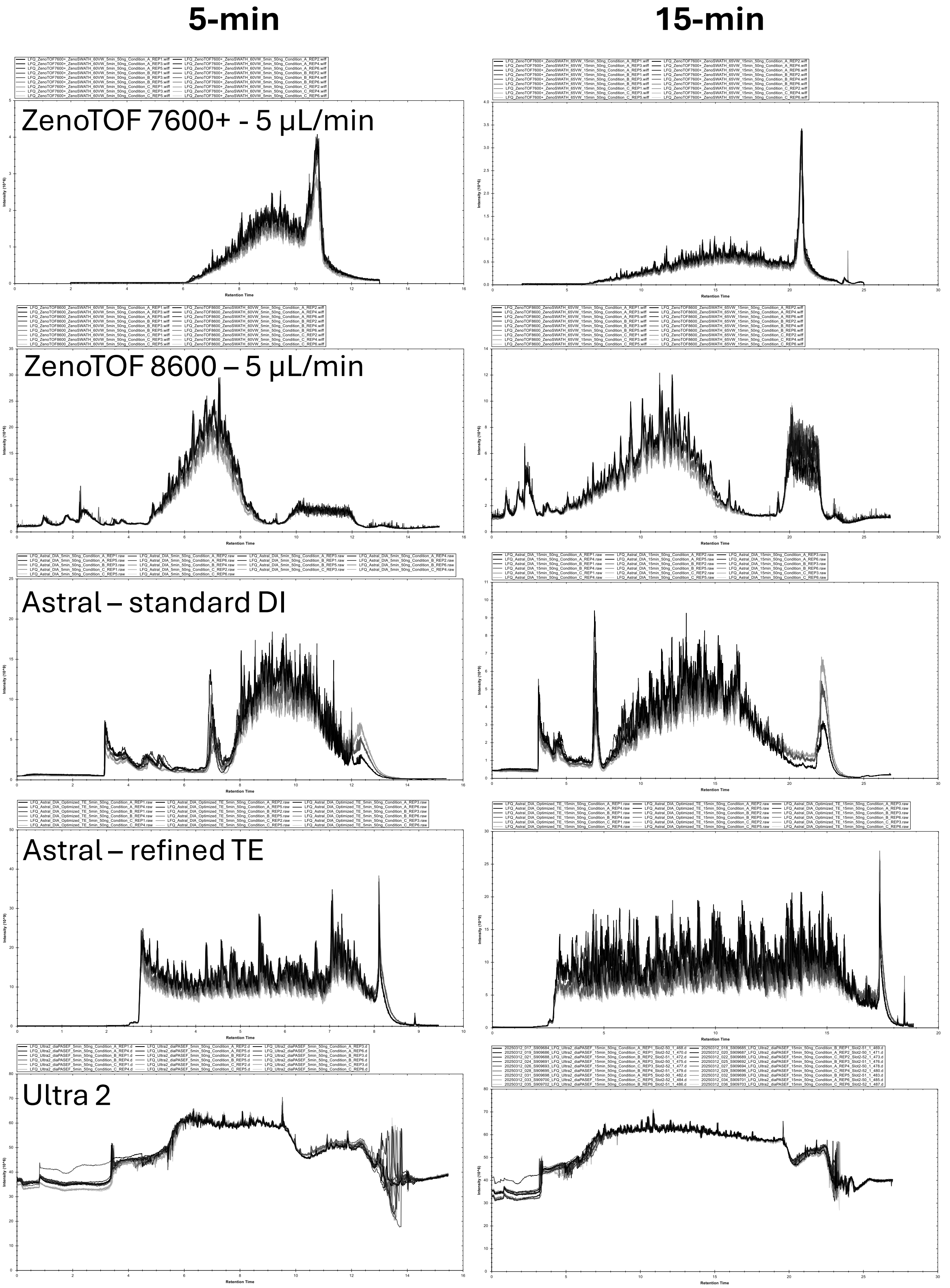

Supplementary Figure 2. TIC overlay plots (all 18 replicates of 50 ng injections) from top to bottom: a) ZenoTOF 7600+, b) ZenoTOF 8600, c) Orbitrap Astral (standard conditions), d) Orbitrap Astral (refined Trap-elute) and e) timsTOF Ultra 2. Plots were generated using Spectronaut.

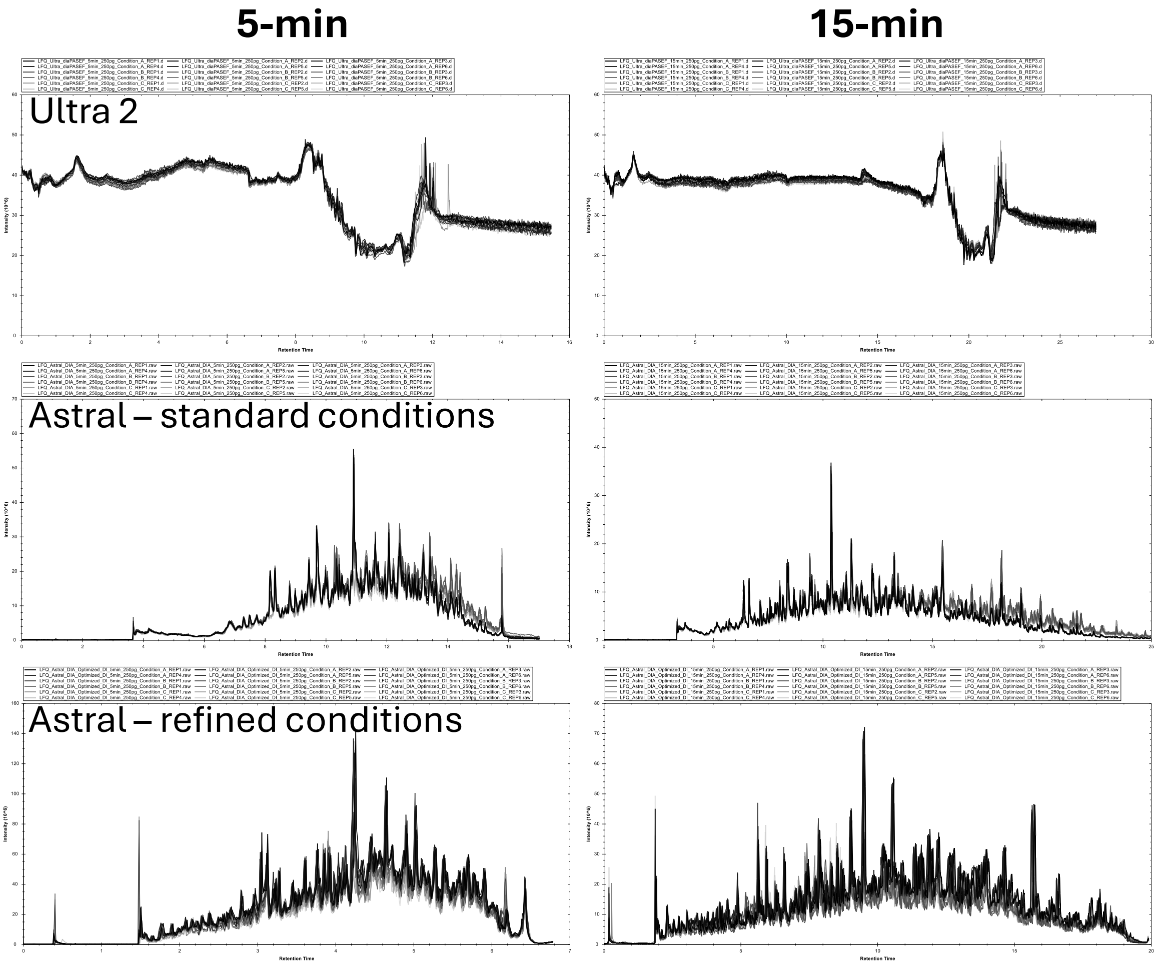

**Supplementary Figure 3**. TIC overlay plots (all 18 replicates) from top to bottom for the 250 pg injections: a) Ultra 2 b) Orbitrap Astral (standard condition) and c) Orbitrap Astral (refined condition). Plots were generated using Spectronaut.

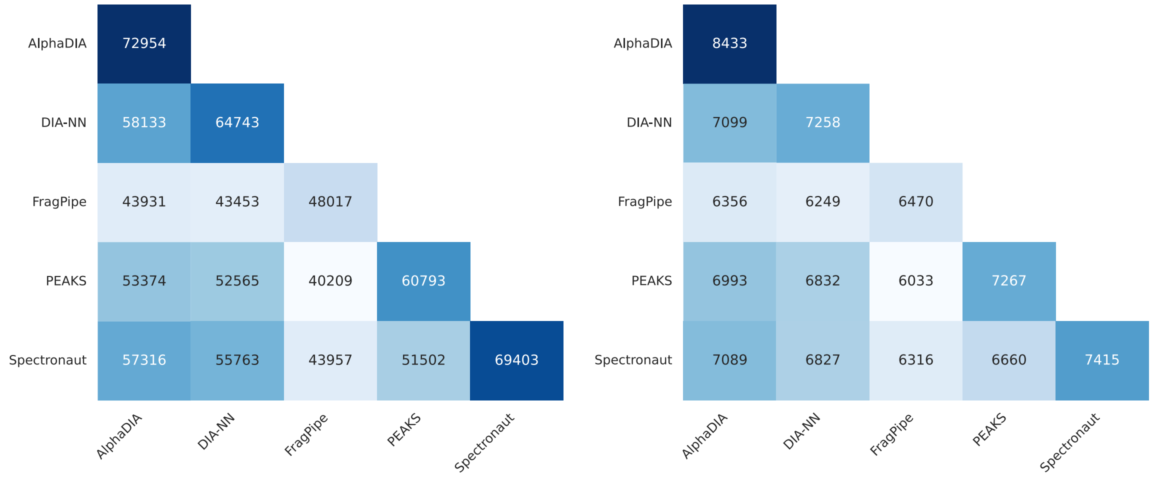

**Supplementary Figure 4.** Overlap in quantified precursor (left) and proteins (right) across different DIA data analysis tools (using a predicted spectral library), for which abundance is not zero, across all replicates of Condition A, B and C (n = 18) measured using the 15-min LC gradient, on the standardized

Orbitrap Astral dataset. For the protein overlap (right), proteins were reassigned independently from the data analysis tool's output using the razor protein inference approach.

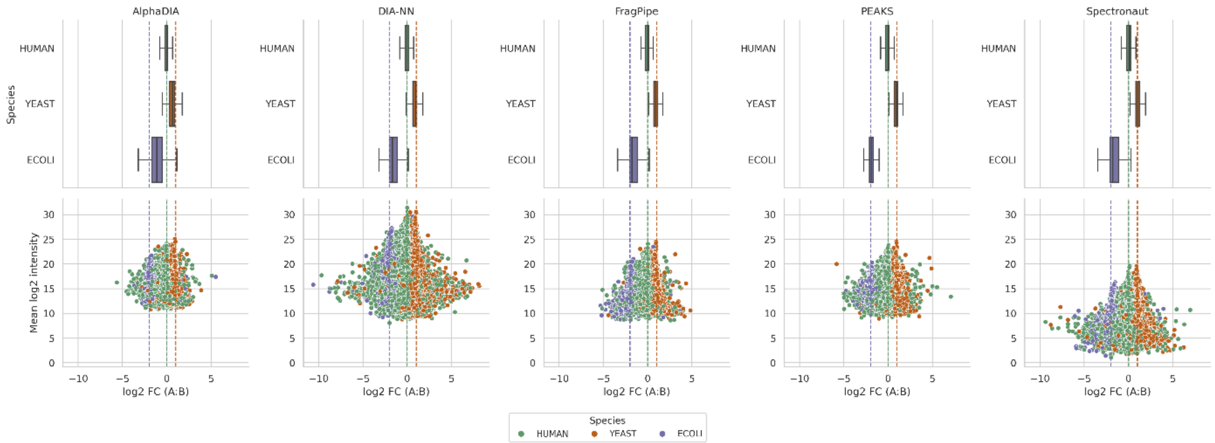

**Supplementary Figure 5.** Comparison of quantitative performance across DIA data analysis tools for the 15-min LC gradient, standardized setup Orbitrap Astral dataset. Top panels: boxplots of $log_2$FC (A:B) for each tool (AlphaDIA, DIA-NN, FragPipe-DIA-NN, PEAKS and Spectronaut). Outliers, defined as data points beyond 1.5 times the interquartile range (IQR) from the quartiles, are not shown. Coloured dashed lines indicate the expected $log_2$FC for each species. Bottom panels: corresponding MA plots, illustrating the relationship between mean $log_2$ intensity (y-axis) and measured $log_2$FC (x-axis) for each precursor.

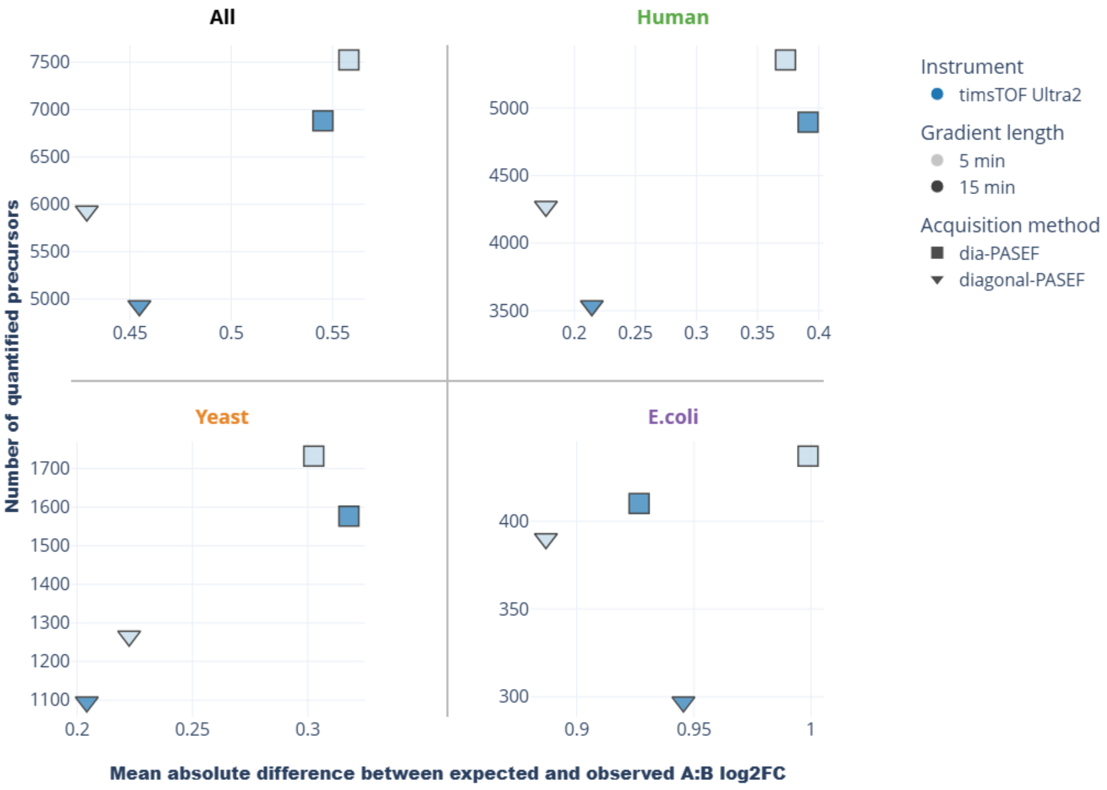

Supplementary Figure 6. Comparison of label-free quantification performance between the Ultra 2 250 pg datasets. These scatterplots show the relationship between the number of quantified precursor ions and the mean quantification error, calculated as the mean absolute deviation to the expected log2FC between condition A and B, for separate species. Metrics are calculated only for precursors identified in all 6 replicates. For the "All" subplot, the average was taken of the mean quantification errors of each species. Precursor ion identification and quantification was performed using DIA-NN (see Supplementary Methods).

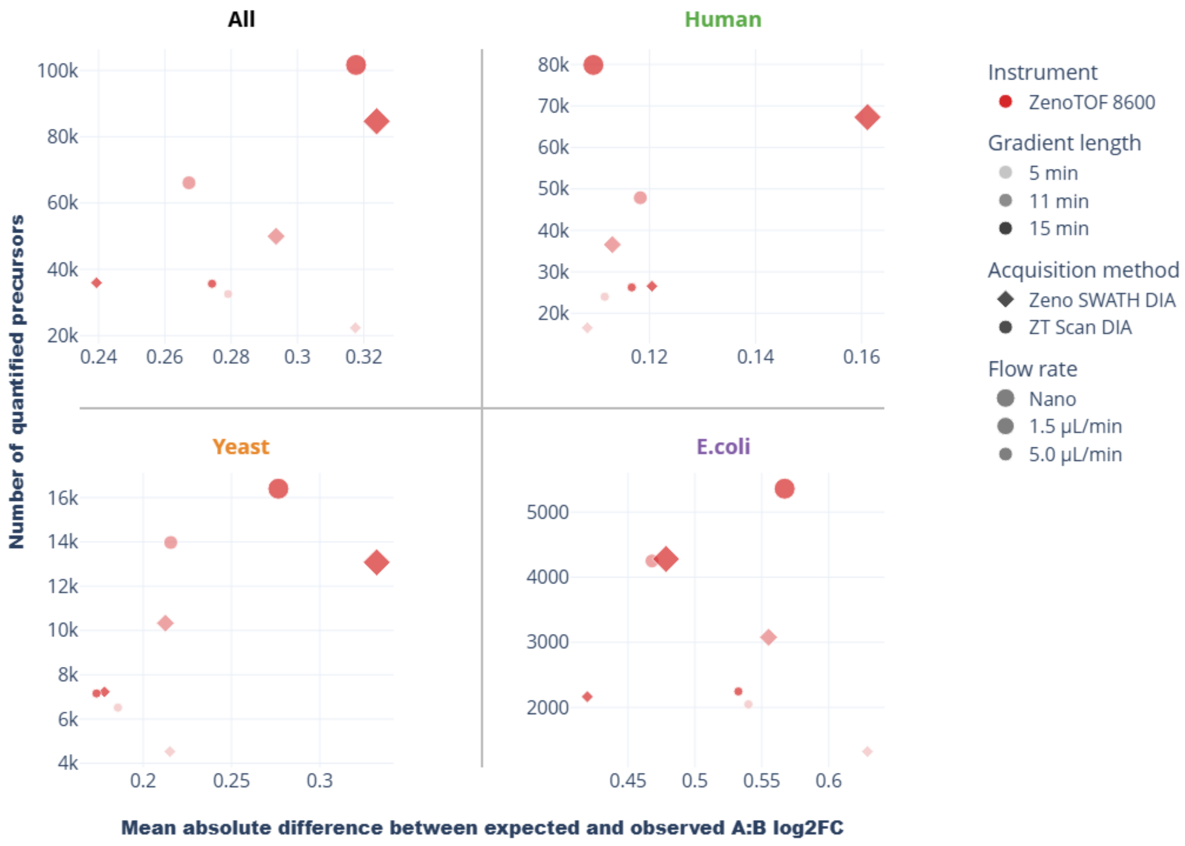

Supplementary Figure 7. Comparison of label-free quantification performance between the ZenoTOF 8600 datasets. These scatterplots show the relationship between the number of quantified precursor ions and the mean quantification error, calculated as the mean absolute deviation to the expected log2FC between condition A and B, for separate species. Metrics are calculated only for precursors identified in all 6 replicates. For the "All" subplot, the average was taken of the mean quantification errors of each species. Precursor ion identification and quantification was performed using DIA-NN (see Supplementary Methods).

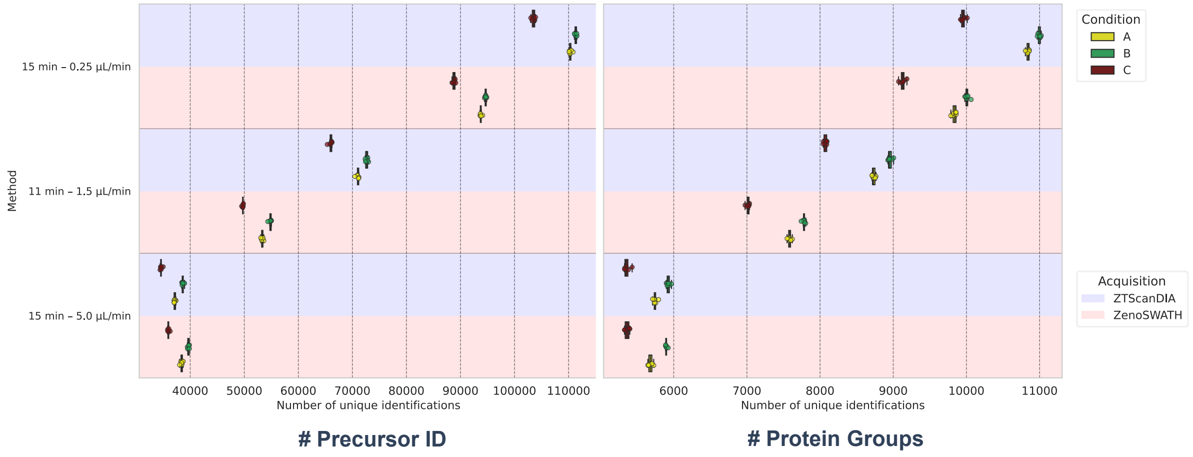

Supplementary Figure 8. The use of different flow rates with the Optiflow Pro source on the ZenoTOF 8600 showed an inverse relationship between flow rate and identification efficiency (based on DIA-NN v2.1.0 output) across all conditions, with lower flow rates yielding higher numbers of identified precursors and protein groups.

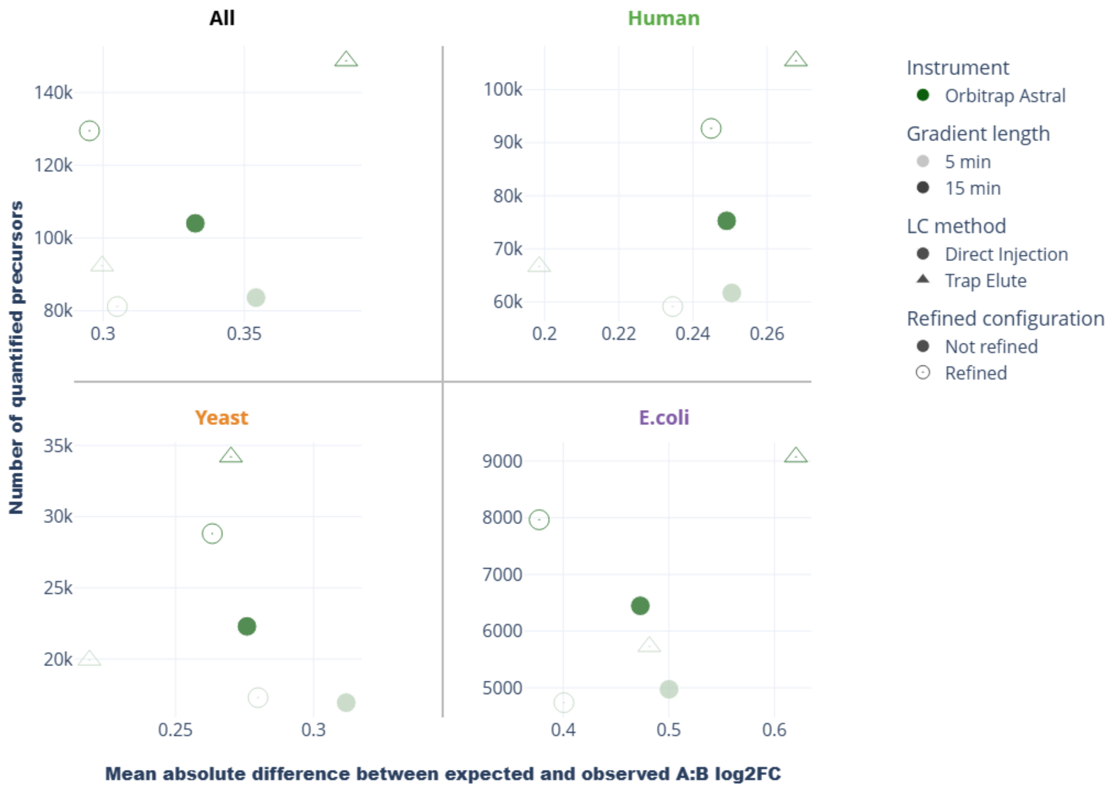

Supplementary Figure 9. Comparison of label-free quantification performance between the different standardized and refined Orbitrap Astral 50 ng datasets. These scatterplots show the relationship between the number of quantified precursor ions and the mean quantification error, calculated as the mean absolute deviation to the expected log2FC between condition A and B, for separate species. Metrics are calculated only for precursors identified in all 6 replicates. For the “All” subplot, the average was taken of the mean quantification errors of each species. Precursor ion identification and quantification was performed using DIA-NN (see Supplementary Methods).

### Supplementary Methods

#### DIA data analysis

##### *General*

All proteomics data analyses were performed in library-free mode, using a FASTA database comprising UniProtKB/Swiss-Prot protein sequences for *Homo sapiens*, *Escher* DIA [illegible]*oli*, and *Saccharomyces cerevisiae*, combined with a universal contaminant protein database [1]. For the figures shown in this manuscript, only the LC-MS/MS runs of the A and B conditions (n = 12) were utilized to generate the results. The precursor *m/z* range used to generate the spectral library was set according to the respective *m/z* range covered by the DIA windows. Carbamidomethylation (C) was used as fixed modification and Acetyl (Protein N-term) and Oxidation (M) were used as variable modifications

##### *DIA-NN*

DIA-NN (v2.1.0) was used to perform precursor ion identification and quantification needed to generate the results shown in Figure 2 and Supplementary Figure 2-7. Trypsin/P was selected as the proteolytic enzyme, allowing up to one missed cleavage, and precursor ion charge states from +1 to +5 were considered. For Orbitrap Astral and ZenoTOF analyses, MS1 and MS2 tolerance was set to 0 to enable automatic optimization. For timsTOF analyses, MS1 and MS2 tolerance was set to 15 ppm, as is suggested by the DIA-NN documentation.

##### *AlphaDIA*

AlphaDIA (v.1.12.1) was used to perform precursor ion identification and quantification needed to generate the results shown in Supplementary Figure 2 and 3 (Orbitrap Astral data). Trypsin/P was selected as the proteolytic enzyme, allowing up to one missed cleavage, and precursor ion charge states from +1 to +5 were considered. A maximum of one variable modification was allowed per precursor ion. MS1 and MS2 tolerance was set to 0 to enable automatic optimization.

##### *FragPipe*

FragPipe (v23.0) was used to perform precursor ion identification and quantification needed to generate the results shown in Supplementary Figure 2 and 3 (Orbitrap Astral data). The "DIA_SpecLib_Quant" workflow was enabled, which uses MSFragger-DIA (v4.3) to perform direct DIA identification to build the spectral library, after which DIA-NN (v 1.8.2 Beta 8) is used for precursor quantification. Trypsin/P ("stricttrypsin") was selected as the proteolytic enzyme, allowing up to one missed cleavage, and precursor ion charge states from +1 to +5 were considered. A maximum of one variable modification was allowed per precursor ion. MS1 and MS2 tolerances were optimized automatically by DIA-NN.

##### *PEAKS*

PEAKS (v13) was used to perform precursor ion identification and quantification. The Enzyme was set to trypsin/P, the Instrument to ZenoTOF/Orbitrap (Astral)/timsTOF, the Fragment type to HCD for Orbitrap (Astral) and CID for the others, and the Acquisition mode to DIA. Data were processed using the Proteome workflow. Search parameters were configured in the Database search parameters tab, allowing up to one missed cleavage, and precursor ion charge states from +1 to +5 were considered. In the Quantification tab, label-free quantification was selected, and samples were added individually or grouped by condition and no normalization was applied. Within the Report tab, both the Precursor FDR and Protein Group FDR were set to 1%.

#### *Spectronaut*

Spectronaut (v 19.9.250422.62635) was used to perform precursor ion identification and quantification using the DirectDIA+ workflow. Analysis settings were configured according to the default DirectDIA+ parameters. No GO terms or library extensions were enabled. After the search completed, the Report tab was used to generate a BGS Factory Report, which was exported as a "_Report.tsv" file.

### Evaluation of quantification performance

Quantification accuracy is defined as the difference between the empirical $\log_2$ fold $\log_2$FC and the expected $\log_2$FC, as determined from the predefined mixing ratios of the experimental conditions. Firstly, in all figures that gauge quantification performance, all precursors mapping to proteins from multiple species or to contaminant proteins were removed prior to further analysis. For a comparison between conditions A and B, precursor intensities were first $\log_2$-transformed. For each precursor, the mean $\log_2$ intensity was then calculated separately for conditions A and B. The empirical $\log_2$FC was obtained by subtracting the mean $\log_2$ intensity in condition B from that in condition A. Quantification accuracy was subsequently computed as the deviation of this empirical $\log_2$FC from the corresponding expected $\log_2$FC.

For Supplementary Figures 5, 6, and 8, this measure of quantification accuracy was further generalized by calculating the mean deviation between the observed and expected $\log_2$FC values across all precursors.
